# A necroptotic-to-apoptotic signaling axis underlies inflammatory bowel disease

**DOI:** 10.1101/2024.11.13.623307

**Authors:** Jiyi Pang, Aysha H. Al-Ani, Komal M. Patel, Samuel N. Young, Isabella Kong, Jin-jin Chen, Marilou Barrios, James A. Rickard, Siqi Chen, Siavash Foroughi, Wayne Cawthorne, Annette V. Jacobsen, Asha Jois, Ashley L. Weir, Lachlan W. Whitehead, Pradeep Rajasekhar, Christopher R. Horne, Imadh Azeez, Tao Tan, Weiwei Liang, Suresh Sivanesan, Andrew Metz, Ash Patwardhan, Natalie Shea, Guru Iyngkaran, Daniel Schneider, Alexander T. Elford, William Beattie, Finlay Macrae, Gianmaria Liccardi, Henning Walczak, Yuxia Zhang, Oliver M. Sieber, Tim Spelman, Lisa Giulino-Roth, Edwin D. Hawkins, Kelly L. Rogers, Rory Bowden, Sandra E. Nicholson, Kate E. Lawlor, Britt Christensen, Andre L. Samson, James E. Vince, James M. Murphy

**Affiliations:** The Walter and Eliza Hall Institute of Medical Research; Parkville, VIC 3052, Australia; University of Melbourne; Parkville, VIC 3052, Australia; Royal Melbourne Hospital; Parkville, VIC 3052, Australia; Department of Medicine, University of Melbourne; Parkville, VIC 3052, Australia; Pediatric Hematology/Oncology, Weill Cornell Medical College; New York, NY 10065, USA; College of Life Sciences, Nankai University; Tianjin, 300071, China; Royal Children’s Hospital; Parkville, VIC 3052, Australia; Drug Discovery Biology, Monash Institute of Pharmaceutical Sciences, Monash University; Parkville, VIC 3052, Australia; Clinical Research Center for Pediatric Infection and Immunity, and Department of Gastroenterology, Guangzhou Women and Children’s Medical Center, Guangzhou Medical University; Guangzhou, 510623, China; The Third Affiliated Hospital of Zhengzhou University; Zhengzhou, 450052, China; Institute of Biochemistry I, Medical Faculty, University of Cologne, 50931, Cologne, Germany; Centre for Cell Death, Cancer and Inflammation, UCL Cancer Institute, University College London WC1E 6BT, London, United Kingdom; Centre for Innate Immunity and Infectious Diseases, Hudson Institute of Medical Research; Clayton, VIC 3168, Australia; Department of Molecular and Translational Science, Monash University, Clayton, VIC 3168, Australia

## Abstract

Inflammatory bowel disease (IBD) is a chronic condition caused by altered cytokine signaling, maladaptive immunity, dysbiosis, and intestinal barrier dysfunction. Patients with IBD receive therapy to correct these imbalances and achieve remission. However, most patients relapse, suggesting that pathological mechanisms persist during remission. Here, we show that excess epithelial cell death is an underlying feature of IBD that arises in patients in remission and on advanced therapy. Mechanistically, nascent inflammation reprograms epithelial cells into a macrophage-like state that promotes RIPK1-independent necroptotic signaling, then triggers iNOS-mediated mitochondrial apoptosis of absorptive epithelial cells and PUMA-mediated intestinal stem cell death. These findings reveal aberrant epithelial cell death signaling as a hallmark of IBD that occurs early in mucosal lesion development and persists despite current therapeutic approaches.

**One-Sentence Summary:** Epithelial cell death is dysregulated in patients with inflammatory bowel disease.

## Introduction

Inflammatory Bowel Disease (IBD) is an umbrella term for chronic and progressive inflammatory disorders that affect the gastrointestinal tract. Crohn’s Disease (CD) and ulcerative colitis (UC) are the main subtypes of IBD which are characterized by alternating periods of disease activity and quiescence, leading to chronic bowel damage and reduced quality of life. By 2030, it is estimated that 1% of the population in Western countries will be affected by IBD, with incidence rising sharply in other industrialized nations (*1*). As the causes of IBD are multifactorial and poorly defined (*2*), the objective of clinical intervention is not curative, but instead to dampen intestinal inflammation using long-term immunosuppressive therapy (*3-5*). Even on advanced therapies, such as Tumor Necrosis Factor (TNF) inhibitors, most patients will experience intermittent flares that necessitate an escalation in treatment, with only 13-34% of patients achieving deep remission as defined by the absence of histological inflammation (*6, 7*). Thus, it is widely believed that underlying drivers of disease persist during remission despite advanced therapies. Identifying these refractory features of IBD is of paramount importance.

Programmed cell death mediates the orderly removal or recycling of tissue matter, which is essential for diverse processes including host defense against pathogens and the prevention of cancer (*8*). Intrinsic apoptosis requires BAX-/BAK-dependent mitochondrial permeabilization, while extrinsic apoptosis relies upon the activation of the initiator caspase, caspase-8 (*8*). Both types of apoptosis culminate in the activation of effector caspases, caspase-3, -6 and -7, that cleave substrates to disassemble the cell (*8*). Necroptosis is an alternate form of programmed cell death that is typically activated when the apoptotic machinery is impaired (*9-13*). Necroptotic signaling causes Receptor interacting serine/threonine kinase (RIPK)-1, RIPK3, Z-nucleic acid binding protein 1 (ZBP1) and TIR domain-containing adapter molecule 1 (TRIF) to variably form a cytosolic scaffold called the necrosome, which in turn promotes the phosphorylation and activation of RIPK3 and Mixed lineage kinase domain-like (MLKL) that can kill cells by necroptosis (*14-19*). Unlike apoptosis, where intracellular constituents are degraded by caspases to limit immunoreactivity, necroptosis involves MLKL-mediated membrane lysis that promotes inflammation via the release of cell contents (*20*).

Since inflammation is a potent inducer of cell death, increased apoptosis and necroptosis have long been linked to IBD (*21-29*). Indeed, the inflammatory cytokines TNF and interferons (IFN) are established therapeutic targets for the IBD field (*30, 31*) and are well-recognized triggers of cell death signaling (*32-35*). Despite this understanding, very few studies have used definitive markers (e.g. cleaved caspase-3) to detect apoptosis (*36-38*) or necroptosis in individuals with IBD (*24, 26, 39-41*). None of these prior studies examined patients on advanced therapies, nor have they profiled multiple cell death modalities. This is a profound gap in knowledge given that inhibitors of RIPK1-mediated cell death are currently being trialed as prospective therapies for UC (*42-44*). Accordingly, here we define the prevalence and mechanisms of apoptotic and necroptotic signaling in adults with IBD within the context of advanced therapies.

## Results

To clarify the relationship between cell death and mucosal lesions in IBD, serial biopsies of non-inflamed, marginally inflamed, and inflamed intestinal tissue were collected from patients with CD or UC (Fig. 1A and Table S1). When patients with IBD presented with no signs of endoscopic inflammation, biopsies were taken from sites that had been classified as non-inflamed, marginally inflamed or inflamed during their prior examinations. As a non-IBD comparator, intestinal biopsies were collected from adults undergoing colonoscopy for non-inflammatory conditions or to exclude malignancy. Over 900 paired biopsies were collected from 80 patients, with even sampling of the main anatomical sites from the ileum to the rectum (Fig. 1B). At the time of endoscopy, the majority of patients with IBD were in remission or had mild disease activity according to clinical (fig. S1A), endoscopic (fig. S1B), and histologic indices (Fig. 1C and fig. S1C). The demographic and clinical characteristics of our study cohort were typical of patients with IBD in Australia (*45*) and in other Western countries, with the exception that a high proportion (61.5%) were on biologic or small molecule therapy (Fig. 1D; collectively referred to as advanced therapy). Accordingly, the molecular events studied herein represent changes that arise during quiescent IBD and despite advanced therapy.

**Fig. 1.**
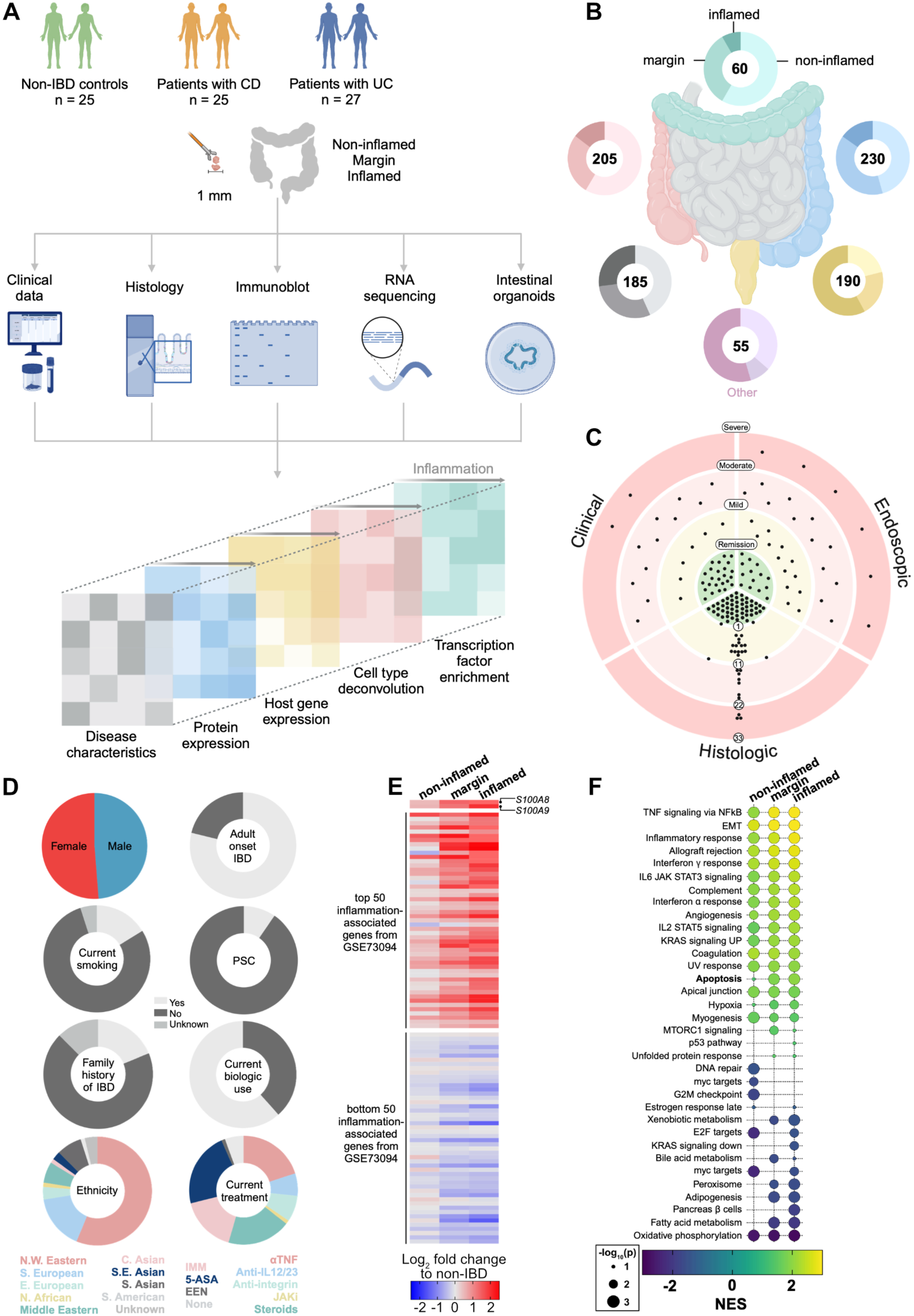
A prospective study to define the role of cell death in IBD. **(A)** Study design. **(B)** The anatomical origin of the biopsies used in this study. **(C)** Disease activity of the cohort at recruitment. Each dot represents one patient with IBD. Clinical activity measured by the Harvey-Bradshaw Index (CD) or the Simple Clinical Colitis Activity Index (UC). Endoscopic activity measured by the Simple Endoscopic Score (CD) or the Mayo subscore (UC). Histopathologic activity measured by the Robarts Histopathology Index (RHI; all cases). **(D)** Demographic or clinical features of cohort at recruitment. Primary sclerosing cholangitis (PSC), thiopurines or methotrexate (IMM), mesalamine (5-ASA), exclusive enteral nutrition (EEN), no therapy (None), infliximab or adalimumab (anti-TNF), Ustekinumab (anti-IL12/23), Vedolizumab (anti-anti-α_4_β_7_), tofacitinib (JAKi), corticosteroid (Steroids). See Table S1. (**E**) Heatmap shows the mean expression of selected genes in intestinal biopsies from patients with IBD (n=26 non-inflamed, n=24 margin, n=25 inflamed biopsies), relative to non-IBD patients (n=23). The genes selected were *S100A8/A9* (encodes for Calprotectin), and the top 50 and bottom 50 inflammation-associated mucosal markers from patients with IBD (*46*). **(F)** Gene set enrichment analysis of bulk RNA sequencing data (same biopsies as Panel E). Normalized enrichment score (NES) and the nominal p-value are shown.

Intestinal samples were subjected to histologic, immunoblot, RNA sequencing, and epithelial organoid assays. Results were centralized in a database that included patient demographic and clinical features (Fig. 1A and Table S1). As a quality control measure, we verified that mucosal markers of IBD (*46*), including *S100A8* and *S100A9*, were expressed in an inflammation-dependent manner in our samples (Fig. 1E) and that blinded histopathologic scores correlated with endoscopic inflammation (fig. S1C). Next, we performed gene set enrichment analysis (GSEA) (*47*) to broadly assess which biological pathways were perturbed in IBD. Despite being on targeted therapies, TNF and interferon (IFN) related genes were still prominently dysregulated in our cohort of IBD samples (Fig. 1F). The expression of necroptosis and ferroptosis gene sets was not altered in IBD samples (fig. S1D), however, apoptosis-related genes were dysregulated in an inflammation-dependent manner (Fig. 1F and fig. S1D). The dysregulation of the apoptotic pathway, even in non-inflamed IBD tissue, was hard to reconcile with the fact that apoptosis-related polymorphisms are not heritable risk factors for adult-onset IBD (*48*). Indeed, unlike in graft-versus-host disease where intestinal apoptosis is a hallmark feature (*49*), histopathological assessment failed to detect appreciable levels of apoptotic cells in IBD tissue (fig. S1E). We therefore reasoned that if apoptotic signaling is dysregulated in IBD, then it involves a mechanism that has thus far been overlooked.

### Elevated cell death signaling is a feature of IBD

To formally address whether apoptotic signaling is dysregulated in IBD, we analyzed the expression and activation of the apoptotic pathway in IBD tissue using immunoblotting, where signals from tissue were quantified relative to well-defined signals in the cultured human HT29 cell line (Fig 2A-C). The necroptotic pathway was also profiled because anti-necroptotic agents are prospective therapies for IBD (*42-44*). In total, 41 different markers of apoptotic and necroptotic signaling were measured across our cohort using immunoblotting (average of n=23 patients/marker; Fig. 2B and fig. S2). This analysis showed that increased necroptotic (defined by phosphorylated RIPK3; pRIPK3) and apoptotic signaling (defined by the ratio of cleaved to full-length caspase-3) are prevalent features of IBD (Fig. 2A-2C). Activation of both pathways correlated with intestinal inflammation in patients with UC or CD, and occurred irrespective of treatment class (Fig. 2A-2C).

**Fig. 2.**
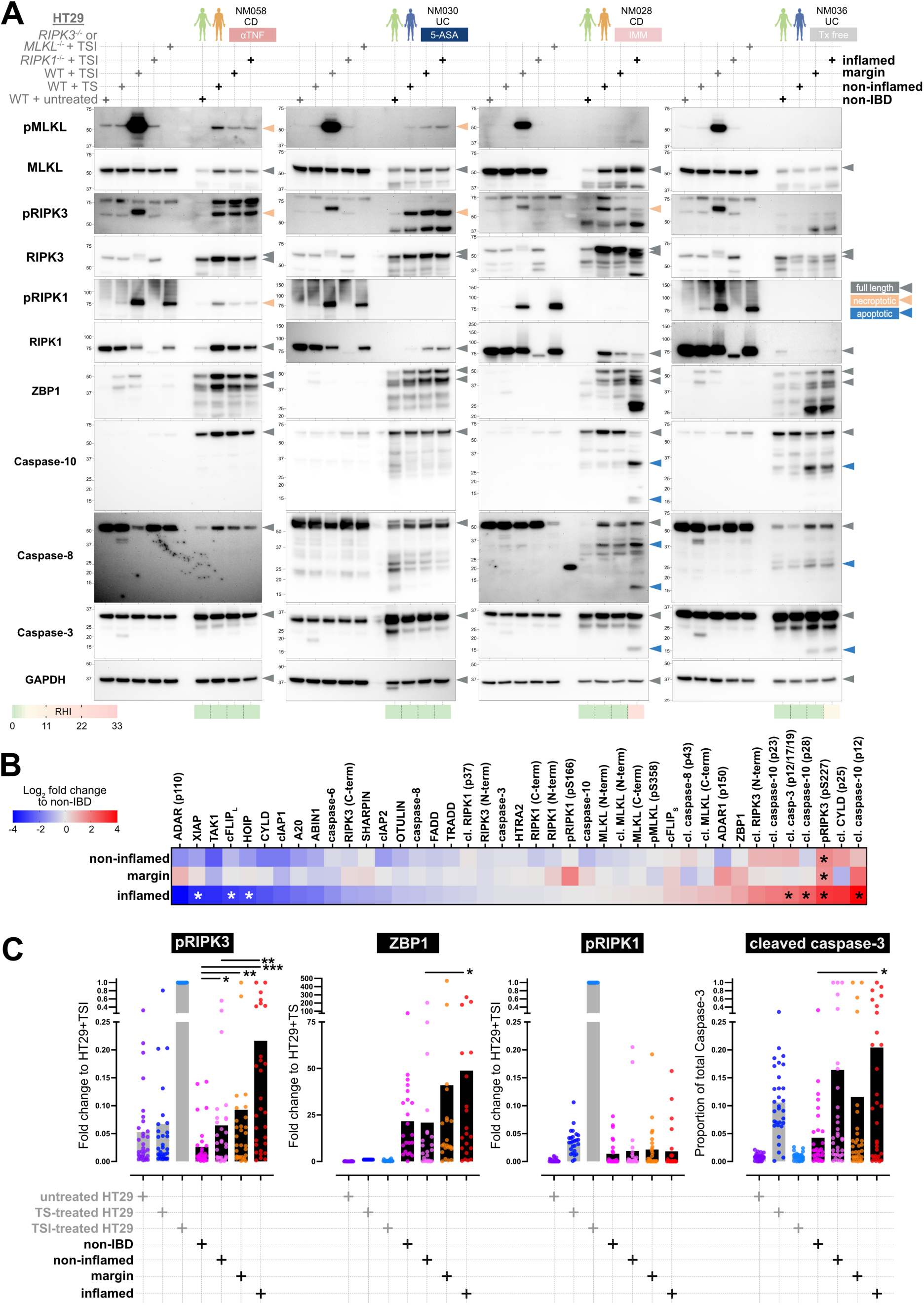
Necroptotic and apoptotic signaling is increased in IBD. **(A)** Immunoblot of lysates from HT29 cells (grey text; left) and intestinal biopsies from patients (black text; right). Apoptotic signaling induced by TNF and Smac mimetic (TS). Necroptotic signaling induced by TS and IDN-6556 (TSI). The fifth lane of each gel contained lysates from TSI-treated *RIPK3^-/-^* or TSI-treated *MLKL^-/-^* cells (see source data for details). The study number (NM), IBD subtype (UC/CD) and treatment for each patient is stipulated. The histopathological score (RHI) of each biopsy site is shown. **(B)** Heatmap shows the expression levels of the indicated proteins or post-translation modifications in biopsies from patients with IBD relative to non-IBD patients. Data are median values (from an average of n=23 biopsies/target/endoscopic grade). *p<0.05 by one-way ANOVA with Geisser-Greenhouse correction. **(C)** Graphs show relative expression levels of the indicated proteins or post-translation modifications in HT29 cells and biopsies. Each dot represents one biopsy. Bars indicate mean values. *p<0.05, **p<0.01, ***p<0.001 by one-way ANOVA with Geisser-Greenhouse correction.

Unexpected traits in necroptotic signaling were also observed. For instance, increased necroptotic signaling was detected in IBD samples, even in ostensibly healthy IBD tissue distal to inflamed tissues, suggesting that necroptotic activation is an early event in disease progression (16 of 31 non-inflamed IBD samples, and 54 of 89 IBD samples in total, had pRIPK3 levels above the non-IBD mean; e.g. patient NM058 in Fig. 2A). While *in vitro* models of RIPK3 activation (*50*) rely on the suppression of key regulators (e.g. caspase-8, cFLIP_L_, HOIP, FADD, OTULIN, IAPs; Fig. 2B and fig. S2A-B) and the co-activation of RIPK1 (Fig. 2C), RIPK3 activation in IBD did not follow this pattern. Instead, RIPK3 activation correlated closely with ZBP1 levels (Fig. 2C). Additionally, there was a subset of patients where increased RIPK3 activation culminated in MLKL phosphorylation, a marker of MLKL activation (21 of 89 IBD samples had both pRIPK3 and pMLKL levels above the non-IBD mean; e.g. patient NM058 Fig. 2A). Furthermore, there was a subset of patients where RIPK3 activation correlated with RIPK1 activation (20 of 79 IBD samples had pRIPK1 and pRIPK3 levels above the non-IBD mean; e.g. patient NM058 Fig. 2A), indicating that patient stratification will likely be important for the therapeutic use of RIPK1 inhibitors (*42-44*). The disconnect between RIPK1, RIPK3, and MLKL activation might reflect immunoblot sensitivity limitations, stochasticity in canonical RIPK signaling, differences in the timing of pathway activation relative to biopsy collection, or the involvement of non-canonical RIPK3 signaling.

The relationship between intestinal inflammation and apoptosis was distinct. For instance, increased apoptotic signaling was observed in histologically inflamed IBD tissue (16 of 30 inflamed IBD samples, and 38 of 93 IBD samples in total, had cleaved caspase-3 levels above the non-IBD mean; e.g. patient NM028 and NM036 Fig. 2A), suggesting apoptosis plays a more downstream role in lesion development. Mechanistically, cleavage of caspase-10 and decreased levels of cell death inhibitors (e.g. HOIP, cFLIP_L_, XIAP; fig. S2A-C) closely correlated with active caspase-3 levels. Total and active RIPK3 and MLKL were also preferentially cleaved in samples where caspase-3 was activated, suggesting that inflammation-associated necroptotic and apoptotic signaling occurred in the same cells (fig. S2D). Overall, 74 of 93 IBD samples had levels of pRIPK3 and/or cleaved caspase-3 above the non-IBD mean, showing that elevated cell death signaling is a highly prevalent feature of IBD.

### Inflammation-induced transcription rewires epithelial cell death signaling

Bulk RNA sequencing data from our cohort of intestinal biopsies were interrogated to further explore how inflammation alters cell death signaling. Remarkably, from an expansive list of necroptotic and apoptotic genes, only 6 were significantly upregulated in an inflammation-dependent manner: *NOS2*, *MLKL*, *PDK1*, *ZBP1*, *BCL2A1* and *NFKB2* (Fig. 3A and fig. S3A-B). *BCL2L10* was the only significantly down-regulated gene in this list (Fig. 3A and fig. S3A-B). Meta-analysis of 26 prior studies confirmed that intestinal expression of *NOS2*, *MLKL*, *PDK1*, *ZBP1*, *BCL2A1* and *NFKB2* was elevated in IBD (*51*). These data suggest that changes in the expression of only a few genes may have a profound impact on cell death signaling. We then performed unsupervised hierarchical clustering to identify IBD-associated regulatory networks. Of the 5 networks identified, ‘cluster B’ was notable because it contained *NOS2*, *MLKL*, *PDK1* and *ZBP1* (Fig. 3B). As cluster B was governed by transcription factors and genes linked to M1-macrophages (fig. S3C-D), we theorized that cluster B reflected the infiltration of M1-macrophages into IBD tissue. Contrary to this hypothesis, digital cytometry indicated that M1-macrophages were not enriched in our IBD samples (fig. S3E-F and Table S1). Indeed, only modest increases in neutrophils, dendritic and mast cells were observed in endoscopically inflamed IBD tissue (fig. S3E-F), presumably because most patients in our cohort had well-controlled disease. We therefore queried whether the cluster B signature was expressed by a non-immune resident cell type. Consistent with this logic, analysis of the publicly available single cell RNA sequencing data showed that genes in cluster B were primarily upregulated in absorptive intestinal epithelial cells in patients with IBD (Fig. 3C; (*52*)). Immunohistochemistry confirmed that intraepithelial necroptotic signals (defined by caspase-8 clusters (*41*)) were present in histologically non-inflamed IBD tissue, whereas intraepithelial apoptotic signals (defined by cleaved caspase-3) were associated with histologically inflamed IBD tissue (Fig. 3D). We therefore propose that inflammation triggers a M1 macrophage-like transcriptional state in intestinal epithelial cells. This response involves the upregulation of genes in cluster B that drives necroptotic signaling during nascent lesion development, progressing to apoptotic caspase activation in more advanced lesions (Fig. 3E). This response correlates with epithelial inflammation (defined by increased *LCN2* and *DUOX2*), is not targeted by current IBD therapy (Fig. 3F), and resembles the transcriptional cell death-related response of mouse macrophages to innate immune stimuli (*53, 54*).

**Fig. 3.**
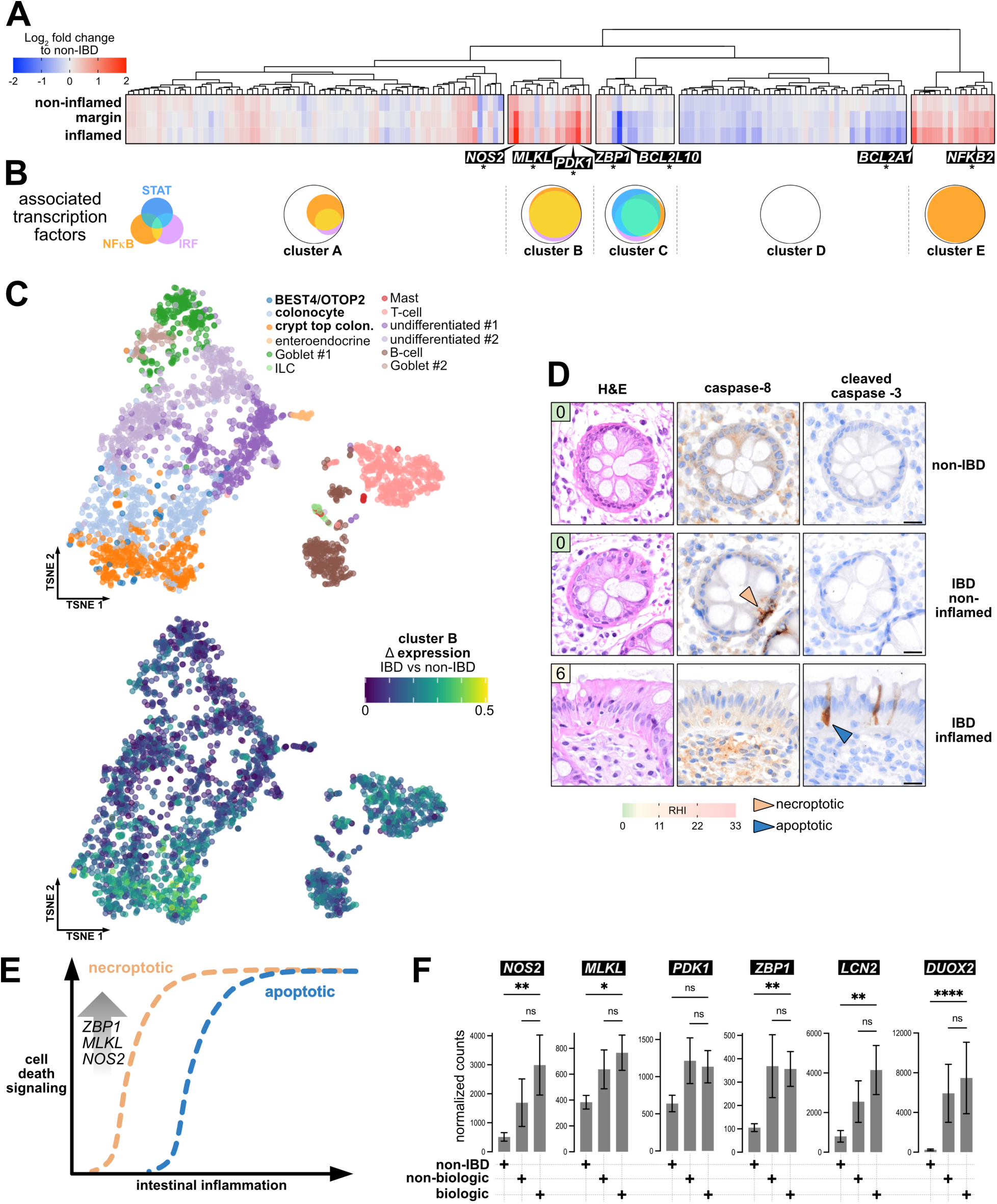
An inflammation-associated transcriptional response promotes epithelial cell death signaling. **(A)** Bulk RNA sequencing of patient samples. Heatmap shows the mean expression of selected genes in intestinal biopsies from patients with IBD (n=26 non-inflamed, n=24 margin, n=25 inflamed biopsies), relative to non-IBD patients (n=23). The selected genes represent 164 key components of the necroptotic and apoptotic pathways (see Methods). Genes that were differentially expressed by a log_2_ fold change of >1 or <-1 and with p<0.05 by multiple testing using the Benjamini–Hochberg method are asterisked. Dendrogram shows the unsupervised hierarchical clustering results used to order genes by similarities in expression and identified 5 distinct groups (clusters A-E). **(B)** Venn diagrams show the main transcription factor families that regulate the genes in clusters A-E. See fig. S3C and Methods. **(C)** Single cell RNA sequencing of intestinal biopsies from **(*123*)**. The t-SNE plots show that increased expression of genes in cluster B is confined to absorptive epithelial cells (bold text) in patients with UC relative to non-IBD patients. Scale bars are 10 units. **(D)** Hematoxylin and eosin (H&E) staining and immunohistochemistry for caspase-8 and cleaved caspase-3 on intestinal biopsies. Intra-epithelial clusters of caspase-8 (orange arrowheads) and apoptotic epithelial cells are indicated (blue arrowheads). Boxed numbers indicate the histopathological score (RHI) of each biopsy site. Scale bars are 2μm. **(E)** Model for how intestinal inflammation triggers an epithelial transcriptional response that promotes necroptotic and apoptotic signaling. **(F)** Transcript levels from the same bulk RNA sequencing data in panel A but grouped according to therapy (non-IBD n=23; non-biologic therapy n=28; biologic therapy n=38 biopsies). Mean ± sem. *p<0.05 and **p<0.01 by one-way ANOVA with Kruskal-Wallis correction. Non-significant (n.s.).

### IFNγ and TNF cooperate to kill intestinal epithelial organoids

The cell death signals observed in IBD tissue correlated with altered TNF and IFN pathway expression (Fig. 1F). Therefore, to further explore how inflammation promotes epithelial cell death, we tested the response of three-dimensional human intestinal organoids to these IBD-associated cytokines, as well as microbial Toll-Like Receptor (TLR) ligands. Organoids of two major epithelial cell types were generated: stem cells and differentiated colonocytes (Fig. 4A). Stem cell organoids were maintained in IntestiCult Organoid Growth Medium (OGM). Differentiated colonocytes were generated in IntestiCult Organoid Differentiation Medium (ODM) supplemented with the Wnt inhibitor IWP2. The phenotype of stem cell- and colonocyte-enriched organoids was confirmed by morphological analysis (Fig. 4A), quantitative PCR (Fig. 4B), immunohistochemistry (fig. S4A), and bulk RNA sequencing (fig. S4B and S4C). Next, we established an unbiased method for quantifying organoid death over time (fig. S4D). Using these approaches, we found that TNF did not cause substantial colonocyte death, while IFNγ induced the killing of ∼40% of colonocytes after 24 hours (Fig. 4C-D and fig. S4E). Interestingly, co-treatment with TNF and IFNγ killed up to 80% of colonocytes within 24 hours (Fig. 4C-D and fig. S4E-F), while no cell death synergy was observed when organoids were co-treated with TNF and IFNα or IFNβ (fig. S4F), by combining IFNγ with TRAIL or FasL (fig. S4G), or by co-treatment with IFNγ and TLR1/2, TLR3 or TLR4 ligands (fig. S4H). Stem cell organoids exhibited similar, albeit less pronounced, death responses to the above stimuli (Fig. 4C-D and fig. S4I). Collectively, these data show that the combination of IFNγ and TNF is an especially potent inducer of epithelial cell death (Video S1).

**Fig. 4.**
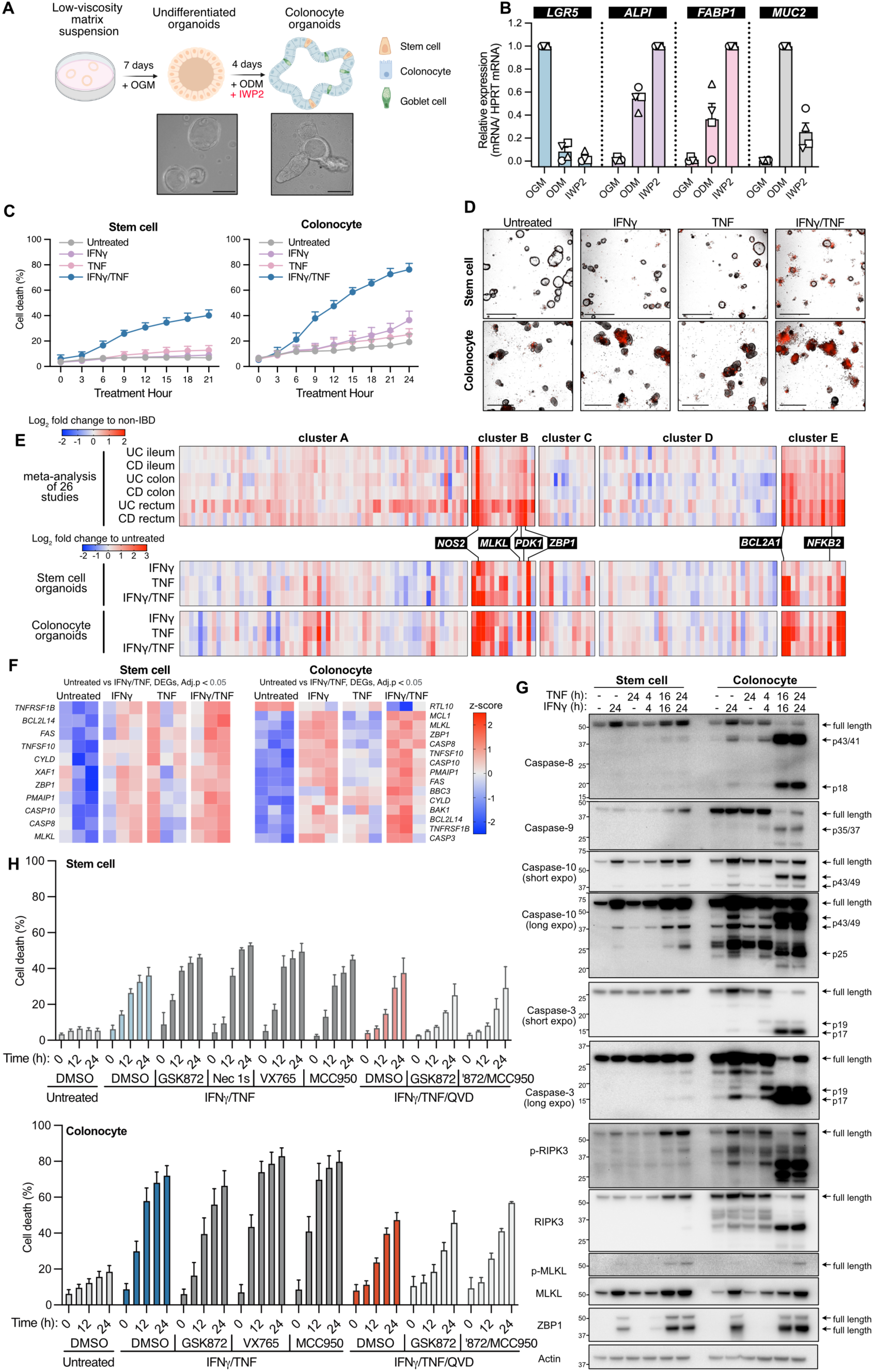
IFNγ synergizes with TNF to trigger intestinal organoid death. **(A)** The intestinal organoid differentiation procedure. Representative micrographs of stem and colonocyte-enriched organoids are shown. Scale bars are 100 µm. **(B)** Expression of stem cell (*LGR5*), goblet cell (*MUC2*) and colonocyte (*ALPI*, *FABP1*) markers was assessed by quantitative polymerase chain reaction (qPCR) in organoids from four donors and normalized to the housekeeping gene *HPRT*, with the group with the highest expression set to 1. **(C, D)** Organoids were treated with IFNγ (50 ng/mL) and/or TNF (50 ng/mL). (C) Percentage death was measured by Cytotox Red-stained organoid area relative to the total organoid area (n = 5 donors, mean ± SEM of 5 independent experiments) using IncuCyte live cell imaging (see fig. S4D). (D) Representative micrographs after 21 hours of treatment; scale bars are 550 µm. **(E)** Heatmaps showing Log_2_ fold change of selected genes in intestinal biopsies from patients with UC or CD, relative to non-IBD patients from published data (*51*) (top) and from for organoids treated with IFNγ and/or TNF for 3.5 hours relative to untreated organoids in this work (bottom). The selected genes, their left-to-right ordering, and the annotated clusters are the same as in Fig. 3A. **(F)** Heatmaps show differential gene expression between untreated organoids and the corresponding organoids treated with IFNγ and/or TNF for 3.5 hours. All entries with an adjusted p<0.05 by empirical Bayes moderated t-statistic and Benjamini–Hochberg multiple test correction. **(G)** Immunoblots of organoids treated with IFNγ and/or TNF as indicated (representative of 3 independent experiments). **(H)** Stem cell organoids were pretreated for 0.5 h with vehicle (DMSO, n=7), RIPK1 inhibitor (necrostatin-1s, 10 μM, n=2), RIPK3 inhibitor (GSK872, 10 μM, n=2), Caspase-1 inhibitor (VX-765, 40 μM, n=3), NLRP3 inhibitor (MCC950, 10 μM, n=2) or the pan-caspase inhibitor (Q-VD-OPh, 40 μM, n=6) alone, or with Q-VD-OPh/GSK872 (n=3) and Q-VD-OPh/ GSK872/ MCC950 (n=3), followed by IFNγ (50 ng/mL) and TNF (50 ng/mL) treatment. Cell death was assessed after 0, 6, 12, 18, 24 hours (n = 5 of different donors, mean ± SEM, pooled from 7 independent experiments). **(I)** Colonocyte organoids were pretreated for 0.5 h with DMSO (n=6), GSK872 (n=3), VX-765 (n=3), MCC950 (n=3) or Q-VD-OPh (n=6) alone, or with Q-VD-OPh/ GSK872 (n=3), Q-VD-OPh/GSK872/ MCC950 (n=3) with the same concentration indicated in (H), followed by IFNγ and TNF treatment. Cell death was assessed after 0, 6, 12, 18, 24 hours (n = 5 of different donors, mean ± SEM, pooled from 6 independent experiments).

### IFNγ transcriptionally reprograms intestinal epithelial cells to promote cell death signaling

Strikingly, the transcriptomic response of organoids to IFNγ and/or TNF (fig. S4J-K) broadly recapitulated gene expression changes observed in the IBD intestine (Fig. 1F). These similarities extended to the apoptotic and necroptotic network, with expression changes in cell death-related genes in inflamed organoids (Fig. 4E) closely mirroring changes in the inflamed IBD samples in our cohort (Fig. 3A) and in a larger cohort of IBD samples (Fig. 4E and (*51*)). Further analysis showed that IFNγ and/or TNF induced the expression of key regulators of apoptosis and necroptosis including death receptors (e.g. *FAS*), caspases (e.g. *CASP8*, *CASP10*), *ZBP1,* and *MLKL* (Fig. 4F and fig. S4K). Immunoblotting of intestinal organoids confirmed that IFNγ increased caspase-8, caspase-10, ZBP1 and MLKL levels, while co-treatment with IFNγ and TNF caused extrinsic (caspase-8, -10), intrinsic (caspase-9), and effector (caspase-3) caspase processing together with activation and cleavage of RIPK3 and MLKL (Fig. 4G). These inflammation-induced apoptotic and necroptotic signals were elevated in colonocytes compared to stem cells (Fig. 4G), consistent with the heightened sensitivity of colonocytes to IFNγ and TNF-induced death (Fig. 4C). Altogether, the treatment of intestinal organoids with IFNγ and TNF emulated the transcriptional and post-translational features of cell death that arise in patients with IBD.

### IFNγ and TNF cause intestinal organoid mitochondrial apoptosis

To define the mode of inflammation-induced epithelial death, we used a panel of inhibitors to block necroptosis (RIPK1 with Nec1s, RIPK3 with GSK’872), pyroptosis (NLRP3 with MCC950, caspase-1 with VX765), death ligands (FASL and TRAIL neutralizing antibodies), or apoptotic caspases (Q-VD-OPh; Fig. 4H and fig. S5A-F). Despite validating their activity in control assays (fig. S5A-B and S5D-F), none of these inhibitors prevented IFNγ and TNF-induced organoid killing (Fig. 4H and fig. S5C). The inability of pyroptosis inhibitors to block IFNγ and TNF-induced organoid death was interesting given the role of Gasdermins in mucosal repair and mucus production (*55, 56*). Even blockade of caspase activity (fig. S5H-I) only delayed the death of IFNγ and TNF-treated organoids, with no additional protection conferred by co-inhibiting necroptotic or pyroptotic signaling (Fig. 4H). We therefore considered whether organoids were dying by mitochondrial apoptosis, where BAX-/BAK-mediated mitochondrial outer membrane permeabilization is sufficient to cause cell death in the absence of caspase activity (*57-59*). This hypothesis stemmed from experiments showing that BH3-mimetics which promote BAX and BAK activation (*60, 61*) still triggered organoid death in the presence of high dose QVD-OPh (with control experiments confirming that QVD-OPh blocked caspase activity; fig. S5D-E and S5G). Accordingly, we engineered organoids to overexpress BCL-2 (fig. S5A-B and S5H-I), the evolutionary conserved inhibitor of BAX and BAK-driven mitochondrial apoptosis (*62*). As expected, BCL-2 overexpression prevented the killing of intestinal organoids by the BH3-mimetic, A-1331852 (targets BCL-XL (*63*)), consistent with cell death occurring via mitochondrial apoptosis (Fig. 5A-C). Similarly, overexpression of BCL-2 prevented IFNγ and TNF-induced death and caspase processing in organoids (Fig. 5A-C). Importantly, the BCL-2 antagonist, ABT199 (*64*), restored IFNγ and TNF-induced killing of BCL-2-overexpressing organoids (Fig. 5B-C). Altogether, IFNγ and TNF triggered both necroptotic and apoptotic signaling in intestinal organoids, however, the activation of RIPK1/3 were not the drivers of epithelial cell death. Instead, mitochondrial apoptosis was the dominant mechanism by which IFNγ and TNF induced both intestinal stem cell and colonocyte death.

**Fig. 5.**
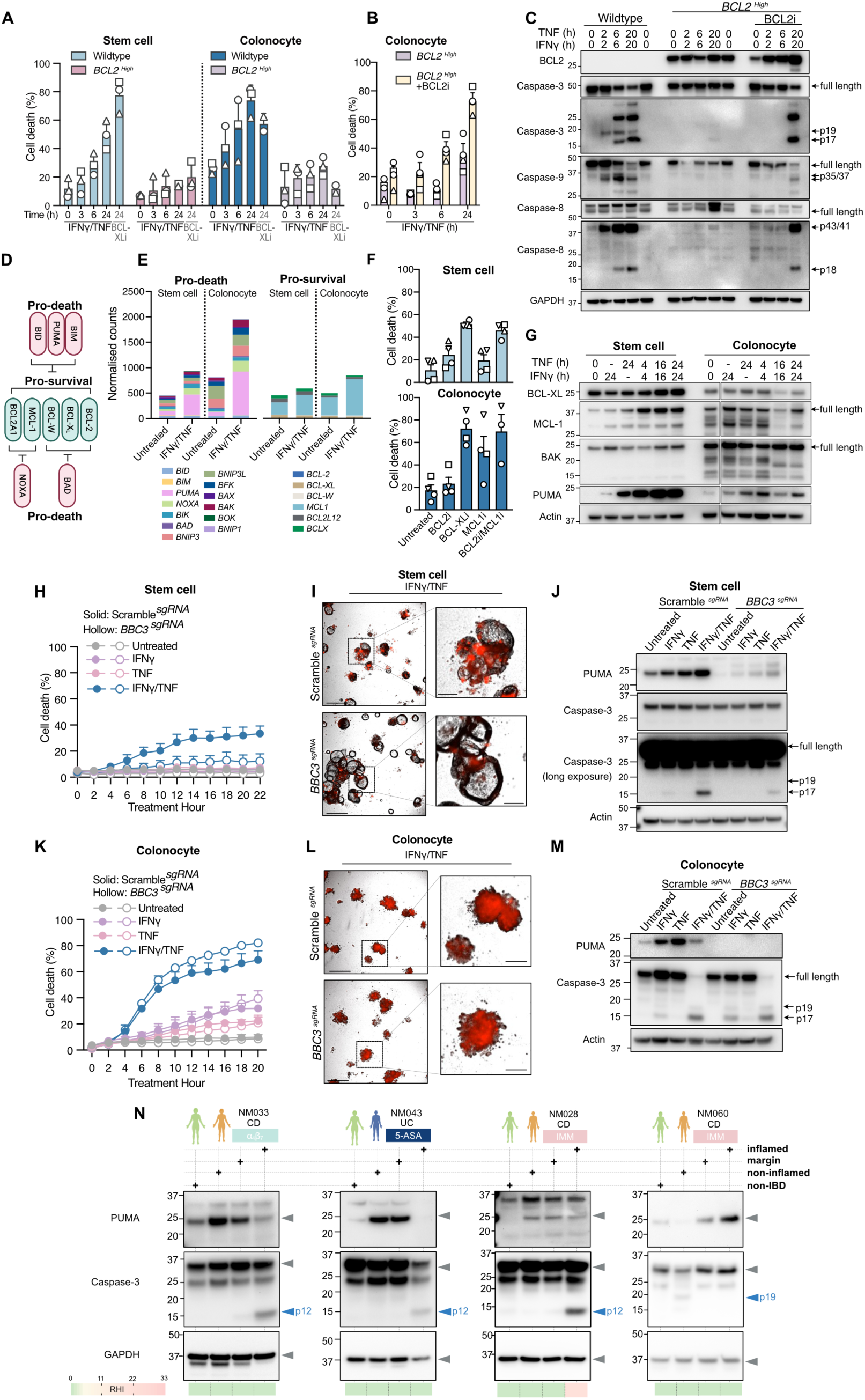
IFNγ and TNF induce mitochondrial apoptosis by upregulating PUMA in intestinal stem cells. **(A,B)** Wildtype control and BCL2-overexpressing organoids were treated with IFNγ and/or TNF and cell death assessed using Cytox Red uptake by IncuCyte live cell imaging (n=3 different donors, represented by different symbols, mean ± SEM, pooled from 3 independent experiments). The BCL-XL inhibitor, A-1331852 (BCL-XLi, 2 μM) was used as a mitochondrial apoptosis control. **(B)** The BCL-2 inhibitor, ABT199 (BCLi, 5 μM) was added to BCL2-overexpressing organoids 30 minutes before IFNγ/TNF treatment and cell death assessed (n=3 different donors, symbols, mean ± SEM, pooled from 3 independent experiments). **(C)** Immunoblot of wildtype and BCL2-overexpressing colonocyte organoids stimulated with IFNγ and TNF with or without ABT199 (5 μM; representative of 3 independent experiments). **(D)** Schematic showing interactions between representative BCL-2 pro-survival family members and pro-death members. **(E)** Sum of normalized transcript counts of pro-apoptotic genes (left) or pro-survival genes (right) from untreated and IFNγ/TNF-treated organoids from 3’ mRNA sequencing (n = 3 different donors). **(F)** Stem cell organoids (n= 4 donors, symbols) or colonocyte organoids (n= 3 donors, symbols) were treated with BH3 mimetics for 24 hours: ABT-199 (2 μM), A-1331852 (2 μM) or S63845 (MCL1i, 5 μM) and cell death assessed (mean ± SEM, pooled from 4 independent experiments). **(G)** Immunoblot analysis of organoids treated with IFNγ and/or TNF (representative of 3 independent experiments). **(H, I)** CRISPR/Cas9 targeted *BBC3* (*BBC3*^sgRNA^) and control (*Scramble*^sgRNA^) stem cell organoids were treated with IFNγ and/or TNF and cell death assessed (n=3 different donors, mean ± SEM, pooled from 3 independent experiments). (I) Representative IncuCyte images of stem cell organoids after 21 hours of IFNγ/TNF treatment. Scale bars are 400 µm (left) and 200 µm (right). **(J)** Immunoblot of *BBC3*^sgRNA^ and *Scramble*^sgRNA^ stem cell organoids treated with IFNγ and/or TNF for 24 hours (representative of 3 independent experiments). (**K, L, M**) Colonocyte organoids were treated and analyzed as indicated in Panels H, I, and J. **(N)** Representative immunoblots showing protein levels of PUMA and cleaved caspase-3 in intestinal biopsies from patients. The patient designation (NM), IBD subtype (UC/CD), and treatment for each patient and histopathological score (RHI) of each biopsy site is shown.

### PUMA drives intestinal stem cell apoptosis

To investigate the mechanisms underlying mitochondrial apoptosis in epithelial cells, we profiled transcript levels of the BCL-2 family – the master regulators of mitochondrial apoptosis (Fig. 5D-F and fig. S5J-K). Colonocyte organoids expressed higher levels of the pro-apoptotic BCL-2 family members than did stem cell organoids under basal conditions, with IFNγ and TNF treatment accentuating this pro-death state in stem cells, and especially in colonocytes (Fig. 5E and fig. S5J). Analysis of publicly available single cell RNA sequencing data confirmed that colonocytes adopted a pro-apoptotic state more than undifferentiated epithelial cells in patients with IBD (fig. S5K; (*52*)). Moreover, gene expression changes within the BCL-2 family were consistent with the capacity of IFNγ and TNF to kill organoids, and aligned with the heightened vulnerability of colonocytes to this insult (Fig. 4C).

To identify proteins that were critical for restraining epithelial apoptosis (Fig. 5D), we treated organoids with inhibitors of BCL-2, BCL-XL and MCL-1; factors that negate BAX and BAK activation. This analysis showed that BCL-XL, but not BCL-2, was essential for preventing stem cell and colonocyte death, with colonocytes also relying on MCL-1 for survival (Fig. 5F). Notably, the pro-apoptotic protein PUMA, which antagonizes BCL-XL and MCL-1, was upregulated in organoids upon IFNγ and TNF treatment, particularly in stem cells (Fig. 5E and 5G). We therefore speculated that PUMA may be important for IFNγ and TNF-induced intestinal cell death. To address this hypothesis, we used CRISPR/Cas9 to delete *BBC3* (encoding PUMA) from organoids. PUMA deletion protected stem cells, but not colonocytes, from IFNγ and TNF-induced caspase-3 processing and death (Fig. 5H-M). These data suggest that inflammation can damage the intestinal stem cell niche by increasing PUMA expression. In support of this concept, a subset of patients with IBD (4 of 11 patients tested) displayed elevated PUMA levels prior to caspase-3 cleavage, implicating PUMA as an initiator of apoptosis in these cases (Fig. 5N and fig. S5L).

### Increased iNOS expression licenses colonocyte death and is upregulated in IBD

As the mechanism of inflammation-induced colonocyte apoptosis was unknown, we questioned whether iNOS, a potent inducer of mitochondrial apoptosis in mouse macrophages (*53, 54*), was also important for the death of inflamed human colonocytes. Notably, despite being a top upregulated gene in IBD tissue (Fig. 6A and fig. S2B), the role of *NOS2* (encoding iNOS) in human biology remains obscure because many human cells, including human macrophages, lack the ability to efficiently express iNOS (*65, 66*). Our hypothesis was prompted by the strong induction of *NOS2* mRNA in IFNγ and TNF-stimulated colonocytes (Fig. 6B), and by immunoblot and immunohistochemistry data showing that iNOS protein levels were highly increased in patients with IBD, particularly within inflamed epithelia (Fig. 6C-F and fig. S6A-C). We confirmed that iNOS protein and nitrite (NO_2_^-^), a marker of nitric oxide production, were induced in colonocytes by IFNγ and TNF treatment (Fig. 6G-H). Deletion of *NOS2* in colonocyte organoids using CRISPR/Cas9 gene-targeting protected against IFNγ and TNF-induced NO_2_^-^ production, apoptotic caspase activation, and cell death (Fig. 6I-L). By comparison, while targeting *NOS2* reduced iNOS and NO_2_^-^ levels in stem cell organoids, it had no impact on IFNγ and TNF killing (Fig. 6M-P), most likely because stem cells produce relatively low levels of iNOS and NO_2_^-^ upon IFNγ and TNF co-treatment (Fig. 6G-H). These data support a model whereby IFNγ and TNF increases iNOS and nitric oxide production to a threshold that can license mitochondrial apoptosis of human intestinal epithelial cells. Collectively, we find that smoldering inflammation in patients with well-controlled IBD rewires intestinal epithelial cells to promote necroptotic signaling and two distinct mechanisms of mitochondrial apoptosis (fig. S6D).

**Fig. 6.**
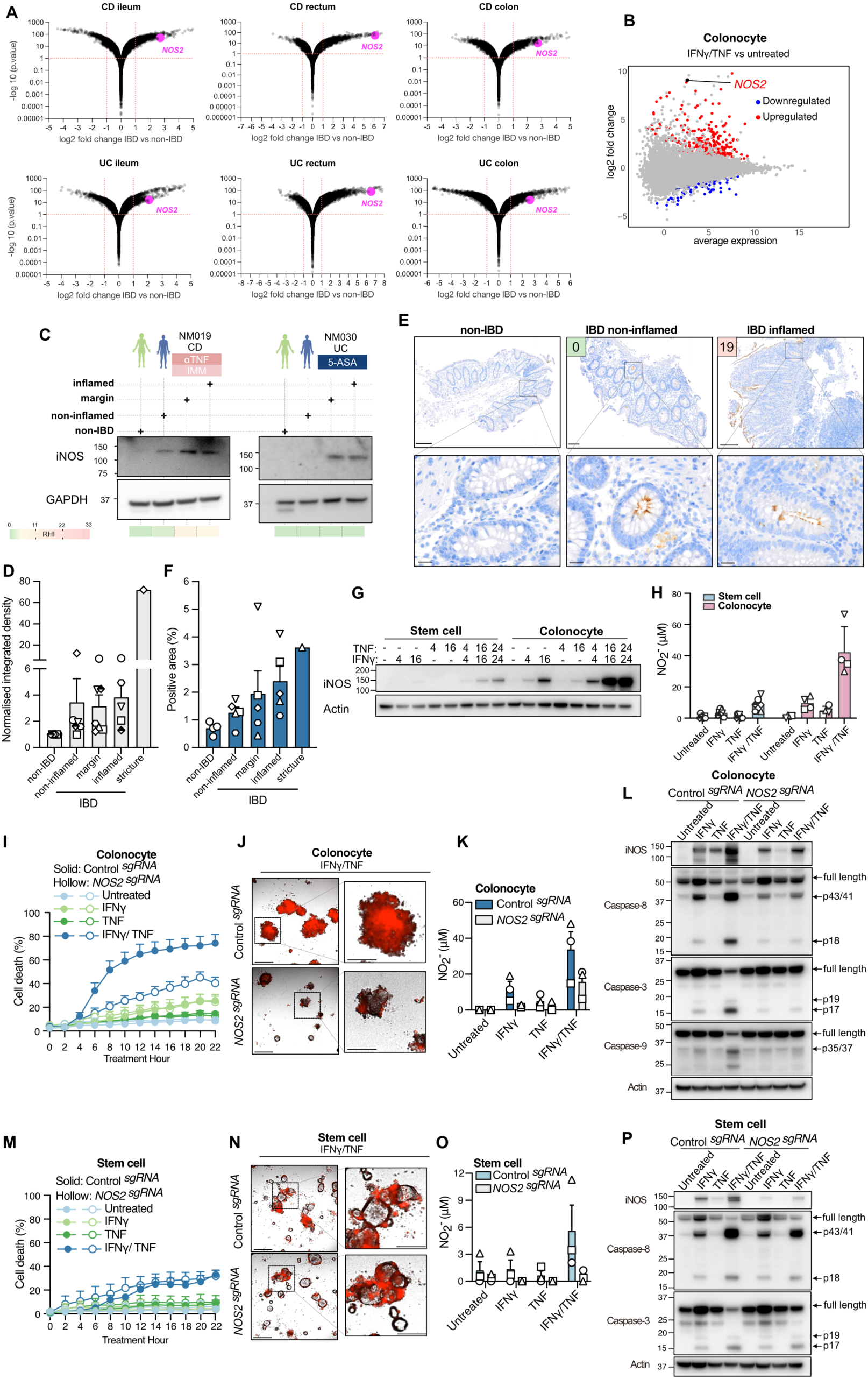
Increased iNOS is detected in IBD tissue and is required for IFNγ/TNF-induced colonocyte organoid killing. **(A)** Increased *NOS2* in patients with IBD relative to control patients. Data combined from 26 studies (*51*). **(B)** Volcano plot showing DEGs and *NOS2* levels in IFNγ/TNF treated colonocyte organoids compared with untreated organoids. **(C, D)** Representative immunoblots and corresponding quantification of iNOS levels in intestinal biopsies from patients with IBD and non-IBD controls (D, mean ± SEM, n =7 per group). See fig. S6 for additional immunoblots. The patient study number (NM), IBD subtype (UC/CD), treatment and histopathological score (RHI) of each biopsy site is shown. **(E, F)** Representative iNOS immunohistochemistry (E) and quantification (F) in IBD intestinal biopsies (mean ± SEM, n=5 patients with IBD patients and n=4 non-IBD control patients). See fig. S6 for additional example of iNOS staining. Scale bars are 200 µm (top) and 20 µm (bottom). **(G, H)** Stem cell and colonocyte organoids were treated with IFNγ and/or TNF and (G) iNOS expression measured by immunoblot (representative of 3 independent experiments) or (H) nitrite (NO_2_^-^) production after 24 hours quantified (n = 4 donors [symbols], mean ± SEM, pooled from 4 independent experiments). **(I, J, K)** The indicated colonocyte organoids were treated with IFNγ and TNF and cell death assessed (I, n=3 different donors mean ± SEM, pooled from 3 independent experiments). Representative IFNγ/TNF treated intestinal organoid images are shown in (J). Scale bars are 400 µm (left) and 200 µm (right). Nitrite (NO_2_^-^) production was measured after 24 hours (K, n=3 different donors mean ± SEM, pooled from 3 independent experiments). **(L)** Immunoblot analysis of colonocyte organoids treated with IFNγ and TNF for 24 hours. Representative of 3 independent experiments. **(M, N, O, P)** Stem cell organoids were treated and analyzed as in **(I, J, K, L)**.

## Discussion

Dysregulated cell death has long been linked to IBD (*23*). Until now, the most definitive evidence linking dysregulated cell death to IBD are cases of early-onset IBD caused by in-born errors in factors that prevent cell death, namely *XIAP* (*67*), *RIPK1* (*68, 69*) and *CASP8* (*70*). Whether the mechanisms of cell death that operate in these rare monogenic cases are similar to those in patients with non-Mendelian IBD is unclear. Our data clearly show that necroptotic and apoptotic signaling in adults with IBD does not rely upon genetic loss of *XIAP*, *RIPK1*, *CASP8* or other related inhibitors of cell death. Instead, we show that nascent inflammation induces the expression of a select group of genes, including *ZBP1* and *MLKL*, and promotes the activation of RIPK3. Notably, because the activation of RIPK3 precedes apoptotic caspase activation during the development of mucosal lesions, necroptotic signaling in IBD is a frontline response to intestinal inflammation, rather than being a secondary response to impaired apoptosis. As an *in vitro* precedent, it has been shown that signaling can be funneled towards a necroptotic outcome simply by changing the levels of a few key pathway components (*71, 72*). Additionally, we show that inflammation also induces the expression of PUMA and iNOS, which can then trigger the death of stem cells and colonocytes. Unexpectedly, while apoptosis is generally considered an immunologically silent process, we find that mitochondrial apoptosis is the prevailing mechanism by which the IBD-associated cytokines, IFNγ and TNF, kill stem cells and colonocytes. Our findings likely explain why co-administration of IFNγ and TNF causes intestinal crypt death in mice (*54*), and attribute function to the expansion of *IFNG^hi^TNF^hi^NOS2^+^*intestinal epithelial cells in patients with CD (*73*). Together, these results highlight inflammation as a potent inducer of epithelial plasticity and cell death signaling in IBD.

An important consideration is whether the cell death signals observed in IBD tissue influence the risk of inflammation-associated colorectal cancer. Notably, mutations in the key cell death-related genes, including *BBC3*, *RIPK3*, *ZBP1* and *MLKL*, are not enriched in colitis-associated neoplastic samples (*74, 75*). Moreover, deletion of *Ripk3* or *Mlkl* does not alter the progression of inflammation-associated colorectal cancer in mice (*76*), suggesting that these cell death effectors do not critically modulate inflammation-driven malignancy. By comparison, somatic mutations in *NOS2* are enriched in patients with colitis-associated neoplasia, leading to the suggestion that iNOS can trigger epithelial apoptosis and thereby guard against malignancy (*74, 75*). Here, we formally show that human iNOS can kill inflamed intestinal epithelia. Given its anti-neoplastic potential, the precise mechanism by which iNOS triggers mitochondrial apoptosis in colonocytes remains of outstanding interest.

Our work has broad clinical ramifications. Durable remission in patients with IBD will likely only be achieved when the molecular drivers of disease are corrected – a goal referred to as molecular healing (*77*). Based on its appearance in histologically normal IBD tissue, we conclude that increased epithelial cell death signaling is an early molecular change in IBD. Given its graded relationship with inflammation, we predict that the extent or type of cell death signaling in IBD will have prognostic value. Furthermore, because dysregulated cell death signaling persists in patients across various treatment regimens and in different states of remission, we propose that it is either a refractory feature of IBD or an aspect that is inefficiently targeted by current therapies. Overall, our findings implicate necroptotic signaling as an upstream event that precedes apoptotic cell death during the progression of IBD, raising the prospect that the therapeutic correction of the necroptotic-to-apoptotic signaling axis may help achieve molecular healing.

## Materials and Methods

### Antibodies

For Fig. 2 and fig. S2, the primary antibodies for immunoblotting were: pMLKL (Rabbit, Ab187091, Abcam), MLKL (Rat, 3H1, in-house (*17*); available from Sigma-Aldrich as MABC604), RIPK1 (Mouse, BD 38/RIP, BD Biosciences), ZBP1 (Rabbit, 5H15L56, Invitrogen), CYLD (Rabbit, CST D6O5O, Cell Signaling Technology), cFLIP (Rabbit, CST D5J1E, Cell Signaling Technology), TRADD (Rabbit, CST 7G8, Cell Signaling Technology), caspase-8 (Rabbit, CST D35G2, Cell Signaling Technology), pRIPK3 (Rabbit, CST D6W2T, Cell Signaling Technology), RIPK3 (Rat, 1H2, In-house (*78*); available from Sigma-Aldrich as MABC1640), GAPDH (Mouse, MAB374, Millipore), pRIPK1 (Rabbit, CST D813A, Cell Signaling Technology), HOIP (Rabbit, CST E6M5B, Cell Signaling Technology), Sharpin (Rabbit, CST D4P5B, Cell Signaling Technology), RIPK3 (Rabbit, CST E1Z1D, Cell Signaling Technology), ABIN (Rabbit, CST 4664S, Cell Signaling Technology), cIAP1 (Rabbit, CST D5G9, Cell Signaling Technology), cIAP2 (Rat, In-house), caspase-8 (Mouse, B.925.8, Invitrogen), RIPK1 (Rabbit, CST D94C12, Cell Signaling Technology), caspase-3 (Rabbit, CST 9662S, Cell Signaling Technology), caspase-10 (Mouse, M058-3, MBL), FADD (Rabbit, Polyclonal, Cell Signaling Technology), ADAR1 (Rabbit, CST E6X9R, Cell Signaling Technology), TAK1 (Rabbit, CST D94D7, Cell Signaling Technology), OTULIN (Rabbit, EPR19841, Abcam), MLKL (Rat, 7G2, In-house (*18*); available from Sigma-Aldrich as MABC1636), A20 (Rabbit, CST D13H3, Cell Signaling Technology), caspase-10 (Rabbit, EPR10890, Abcam), XIAP (Rabbit, EPR22189, Abcam), iNOS (Mouse, MAB950, R&D Systems), PUMA (Rabbit, CST 98672, Cell Signaling Technology), caspase-6 (Rabbit, Polyclonal, Cell Signaling Technology). For Fig. 4-6 immunoblots, the primary antibodies were: caspase-3 (Cell Signaling; 9662), caspase-8 (Cell Signaling; 9746), caspase-9 (Cell Signaling; 9508), caspase-10 (MBL; M059-3), BCL-XL (Cell Signaling; 2764), MCL-1 (Cell Signaling; 5453), PUMA (Cell Signaling; 98672), β-actin (Sigma; A-1798), GAPDH (Millipore, MAB374), iNOS (R&D MAB9502), BCL2 (WEHI; clone Bcl-2-100), BAK (clone 4B5, (*79*)). The secondary antibodies (1:10,000 working concentration) for immunoblot were horseradish peroxidase (HRP)-conjugated goat anti-rat immunoglobulin (Ig) (Southern BioTech Cat#3010-05), HRP-conjugated goat anti-rabbit Ig (Southern BioTech Cat#4010-05), and HRP-conjugated goat anti-mouse Ig (Southern BioTech Cat#1010-05). The primary antibodies for immunohistochemistry were mouse anti-human caspase-8 (clone B.925.8; RRID:AB_10978471; 0.619 g/L Thermo Fisher Scientific Cat# MA5-15226), rabbit anti-cleaved-caspase-3 (Cell Signaling Technology, #9661) and iNOS (R&D MAB9502, 1:200), anti-LGR5 (clone OTI2A2, Thermo Fisher Scientific, TA503316, 1:300) and anti-Intestinal Alkaline Phosphatase Polyclonal Antibody (ALPI, Catalog # PA5-22210, 1:200).

### Bulk RNA sequencing of intestinal biopsies and analysis

Samples were thawed, RNAlater was removed, then tissues were transferred into screw-capped tube pre-filled with 350 µl of RA1 buffer of NucleoSPin RNAXS kit (Macherey-Nagel Cat# SKU: 740902.250). Tissues were homogenized with 10 pcs of 3 mm Acid-Washed Zirconium Beads (OPS diagnostics Cat# BAWZ 3000-300-23) in a Qiagen TissueLyzer II (30 Hz, 5min). Homogenized samples were spun down for 1 minute at 11,000 *xg* to remove tissue debris then RNA was purified using Nucleospin RNAXS column kit as per manufacturer’s instructions without adding a carrier RNA. The purified RNA was quantified using Qubit™ RNA HS Assay kit (Thermo Fisher Scientific Cat#Q32852) and RNA integrity was visualised in high sensitivity RNA ScreenTape (Agilent Cat# 5067-5579) using TapeStation 4200 (Agilent Cat# G2991BA). Ten nanograms of RNA were used for preparing indexed libraries using SMARTer Stranded Total RNA-Seq Pico-Input Mammalian kit v.3 (Takara Bio. Cat# SKU: 634487) using manufacturer’s instructions with a couple of modifications, specifically 3 minutes of fragmentation at 94 °C and 13 cycles of PCR2. Library concentration was quantified by Qubit™ dsDNA Assay kit (Thermo Fisher Cat#Q32851) and library size was determined using D1000 ScreenTape (Agilent Cat# 5067-5582) and visualized in TapeStation 4200 (Agilent Cat# G2991BA). Equimolar amounts of libraries were pooled and loaded for 150-bp paired-end sequencing on one S4 lane of NovaSeq 6000 (Illumina, San Diego, USA) as per manufacturer’s instructions.

The single-end 75 bp were demultiplexed using CASAVA v1.8.2 and Cutadapt (v1.9) was used for read trimming (*80*). The trimmed reads were subsequently mapped to the human genome (grch38) using HISAT2 (*81*). FeatureCounts from the Subread package (version 1.34.7) was used for read counting after which genes <2 counts per million reads (CPM) in at least three samples were excluded from downstream analysis (*82, 83*). Count data were normalized using the trimmed mean of M-values method and differential gene expression analysis was performed using the limma-voom pipeline for pair-wise comparison (limma version 3.40.6) (*82, 84, 85*), or GraphPad Prims v10 for multiple group comparisons. Adjustment for multiple testing was performed per comparison using the false discovery rate method (*86*). Heatmaps of log_2_ fold changes in CPM were generated using GraphPad Prism v10. For pathway perturbation analysis, Gene Set Enrichment Analysis software v2.2.2 was used (*47*) with the Human MSigDB hallmark (H) gene sets (*87*). For expression pattern analysis, unsupervised hierarchical clustering was used to identify subsets of highly correlated genes using the *cluster* R package. Under hierarchical clustering, expression data were first normalized and then the difference or distance between different genes and gene sets were derived to create a distance matrix using the Euclidean distance method. Optimal clusters were determined using the Elbow method, average Silhouette method and/or the Gap statistic method as indicated. The *hclust* function was then used to generate the dendrograms to visually represent the clustering (*88*). For transcription factor enrichment analysis, genes from clusters A-E were submitted to the ChIP-X Enrichment Analysis 3 (ChEA3) tool (*89*) and the 10 most enriched transcription factors considered according to the default settings of ChEA3. Transcription factors were manual assigned to families and conveyed via Venn diagrams generated using the DeepVenn tool (*90*). For digital cell-type deconvolution, the estimation of cell type abundance, and of correlations, in full transcriptome RNA sequencing analysis was performed using CIBERSORTx (*91*). The signature matrix file used for CIBERSORTx analysis was the LM22 file from (*92*). For GSEA, GSEA software v4.3.3 (*47*) was used with all gene sets from GO:BP (gene ontology biological process) of MSigDB (*93*).

### Bulk RNA sequencing of organoids and analysis

Total RNA was isolated from organoids using the ISOLATE II RNA Mini Kit (Bioline, 52073) following the manufacturer’s guidelines. The quality of the extracted RNA was assessed using the Agilent 4200 TapeStation, and RNA samples with RINe values greater than 9 used for further applications. For mRNA sequencing, 100 ng of total RNA was used to prepare 3’ mRNA-sequencing libraries following the QuantSeq 3’ mRNA-Seq Library Prep kit (Lexogen) protocol. The libraries were then sequenced on the NextSeq 500 (Illumina). The single-end 75 bp reads were demultiplexed using Casavav1.8.2 and Cutadapt (v1.9) (*94*) to remove poly A tails as well as adapters, and to remove the low quality reads with a quality score below 20. The reads shorter than 50 bases were discarded to ensure the high quality for the downstream analysis, assessed and validated by using fastQC (*95*). The reads were then aligned to the reference genome GRCh38 using STAR (*96*) and gene expression was quantified at exon level using featureCounts (*82*) to obtain the raw count data. Genes without a current symbol name were removed.

The data filtration, normalization and differential expression analysis was conducted using mastR (*97*). The genes with low counts were filtered out using edgeR::filterByExpr (*98*) prior to the analysis, and raw count data was normalized using TMM (*85*). Differential expression analysis was performed by mastR using a limma-voom-treat (*99*) pipeline with default settings. Differential expression analyses were conducted with a two-factor design to fit a linear model with ‘state’ (cell differentiation state) and ‘treat’ (sample treatment) as the covariates. An empirical Bayes moderated t-statistic was generated and Benjamini–Hochberg multiple testing adjustment was performed to identify statistically significant genes in each comparison (adjusted p-value <0.05). For pathway perturbation analysis, GSEA software v2.2.2 was used (*47*) with the Human MSigDB hallmark (H) gene sets (*87*).

### Cell lines and culturing

HT29 (ATCC, HTB-38), HEK293T (ATCC, CRL-3216), MDA-MB-231 (ATCC, HTB-26) and HepG2 (ATCC, HB-8065) cells were originally sourced from the American Type Culture Collection. The *RIPK1*^−/−^, *RIPK3*^−/−^, *MLKL*^−/−^ HT29 cells have been previously reported (*100-102*). For doxycycline (dox)-inducible-*NOS2* HEK293T cells, pLIX403-hNOS2 (*103*) (a gift from Edward Morgan, Addgene plasmid # 110800; http://n2t.net/addgene:110800; RRID: Addgene_110800) was transiently transfected into HEK293T cells alongside pMDL (packaging), RSV-REV (packaging) and VSVg (envelope) using Lipofectamine 2000 diluted in OptiMEM (Thermo Fisher Scientific) to generate lentiviral particles in DMEM. The cell culture supernatant was collected 48 h later and filtered through a 0.45 mm filter prior to cell transduction. Lentiviral transduction was performed by replacing normal cell culture medium with DMEM containing lentivirus particles for 24 h. Transduced positive cells were selected by puromycin (2ug/mL) for 3 days.

The origin of cell lines was not further verified, although their morphologies and responses to cell death stimuli were consistent with their stated origins. Cell lines were monitored via polymerase chain reaction every ∼6 months to confirm they were mycoplasma-free. Human cell lines were maintained in Dulbecco’s Modified Eagle Medium (Gibco, Life Technologies) supplemented with 8-10 % v/v Fetal Bovine Serum (FBS, Sigma-Aldrich), L-glutamate and 50 U/mL penicillin, 50 μg/mL streptomycin in a humidified incubator at 37 °C and 10 % CO_2_.

### Mouse macrophage generation

Bone marrow-derived macrophages were derived as described (*53*). Bone marrow cells were harvested from femoral and tibial bones and cultured for 6 days in DMEM containing 10% FBS, 50 U/mL penicillin, 50 μg/mL streptomycin, and 15-20% L929 cell-conditioned medium, with an additional 10 mL of 20% L929 cell-conditioned medium added on day 3. At day 6 post-harvest, differentiated BMDMs were replated at 5×10^5^ cells per well in 24-well plates with 500 μL DMEM/FCS and 20% L929 conditioned medium and were used for experiments on day 7.

### Cell line treatment

MDA-MB-231 and HepG2 cells were plated at 2×10^4^ cells per well in 96-well plates with 100 μL DMEM containing 10% FCS. After cells had adhered to the plate, MDA-MB-231 cells were primed with recombinant human IFNγ (50 ng/mL) overnight, followed by TRAIL (50 ng/mL) treatment for 48 hours. TRAIL neutralizing antibody (10 μg/mL) was added 30 minutes prior to TRAIL treatment. HepG2 cells were treated with Actinomycin D, Streptomyces sp (0.5 μg/mL, Sigma-Aldrich, 114666) and FAS ligand (100 ng/mL) for 48 hours, Human Fas Ligand/TNFSF6 Antibody (10 μg/mL) was added 30 minutes prior to FAS ligand treatment. BMDMs was primed with LPS (100 ng/mL) for 3 hours followed by Nigericin (10 mM, Sigma; N7143) treatment. Dox-inducible-*NOS2* HEK293T cells were plated at 1×10^6^ cells per mL in 10-cm dishes, then treated with doxycycline (1 μg/mL, Sigma-Aldrich) overnight.

To make cell culture standards for immunoblotting, HT29 cells were treated in DMEM containing 8% v/v FCS. Media for treatment was supplemented with: 100 ng/mL recombinant human TNF-α-Fc (produced in-house as in (*104*)), 500 nM Smac mimetic/Compound A (provided by Tetralogic Pharmaceuticals as in (*105*)) 5 μM IDN-6556 (provided by Idun Pharmaceuticals). HT29 cells were treated for 7.5 hours.

### Organoid generation and differentiation

Fresh human intestinal samples were collected and kept in DMEM/F-12 with 15 mM HEPES (DMEM/F-12, STEMCELL Technologies, 36254) and 1% w/v Bovine Serum Albumin (BSA, Sigma-Aldrich, A4612) until further processing. Samples were washed 3 times with 5 ml of phosphate-buffered saline (PBS) containing 100 μg/mL Primocin® (Invivogen, ant-pm-05) and 20 μg/mL Gentamicin (Thermo Fisher Scientific, 15710064), then were digested in 3 mM EDTA chelation buffer (Sigma-Aldrich, E5134) containing 100 µM dithiothreitol (Merck, 10197777001) at room temperature for ∼30 minutes, with periodic shaking to facilitate crypt release. Crypts were collected via brief centrifugation and resuspended in DMEM/F-12 containing 1% w/v BSA.

Organoids were then generated and cultured using Low-viscosity matrix suspension culture methods described in (*106*). Briefly, 1 mL of IntestiCult™ Organoid Growth Medium (Human) (OGM, STEMCELL Technologies, 06010) containing 5% Matrigel matrix (Bio-strategy, BDAA354234), 100 U/mL penicillin-streptomycin (Life Technologies, 15140122), 10 μM Rho-kinase inhibitor Y27632 (Stemcell Technologies, 72308) and 20 μg/mL Gentamicin was used to resuspend and plate cyrpts in sterile 24-well non-treated tissue culture plates (Falcon, 351147). Plates were incubated at 37°C with 5% CO_2_. The medium was refreshed every two days by adding 200 μL of fresh OGM into each well. Mature organoids were obtained within 7 to14 days.

For passaging, organoids were collected via centrifugation at 300 x*g* for 3 minutes at 4°C. The pellets were washed once with ice-cold PBS and then incubated with TrypLE Express enzyme (Thermo Fisher Scientific, 12604021) at 37°C for 10 minutes for cell dissociation. Cells were centrifuged at 10,000 x*g* for 30 seconds, and digestion was terminated by adding 1 mL DMEM/F-12 containing 1% BSA. Then cell pellets were mechanically dissociated into single cells using a 26 G needle, and were replated into sterile, non-treated 24-well plates at a density of 1×10^5^ cells per well in a final volume of 1 mL OGM supplemented with 5% Matrigel matrix and 10 μM Rho-kinase inhibitor Y27632. The plates were incubated at 37°C with 5% CO_2_ and medium was refreshed every two days by adding 200 μL of fresh OGM into each well. Mature organoids were obtained within 7 days.

To obtain colonocyte organoids, undifferentiated organoids at day 4-6 were collected, washed twice with ice-cold PBS and resuspended in IntestiCult™ Organoid Differentiation Medium (Human) (ODM, STEMCELL Technologies, 100-0214) supplemented with or without 10 µM IWP-2 (STEMCELL Technologies, #72122) and 5% Matrigel matrix. The resuspended organoids were then replated and incubated at 37°C with 5% CO_2_. After 4 days of differentiation, organoids were used for experiments.

### Organoid stimulation

Organoids were maintained in non-coated 24-well tissue culture plates with OGM until day 7, then were collected via centrifugation at 300 xg for 3 minutes. The pellets were washed once with ice-cold PBS and resuspended in either OGM with 5% Matrigel matrix for stem cell organoid, ODM with 5% Matrigel matrix for goblet organoids, or ODM supplemented with 10 µM IWP-2 and 5% Matrigel matrix for colonocyte organoids. 300-500 organoids were then plated per well in triplicate in non-treated 96-well tissue culture plates (Falcon, 351172) with 50 µL of media. The surrounding wells were filled with PBS, and the plate incubated at 37 °C in a 5% CO_2_ incubator until treatment.

Unless otherwise stated in figure legends, organoids were treated with IFNγ (50 ng/mL, recombinant human, R&D, 285-IF-100), IFNα (100 ng/mL, R&D, 10984-IF), IFNβ (100 U/mL, recombinant human, Prospec, CYT-236), Human Fc-TNF (50 ng/mL, in-house (*107*)), LPS (50 ng/mL, InvivoGen; tlrl-3pelps), Pam-3-CSK4 (500 ng/mL, InvivoGen; tlrl-pms), PolyI:C (10 mg/mL, InvivoGen; tlrl-picw), FAS ligand (10 ng/mL, recombinant human, Peprotech, 310-03H), TRAIL (5 ng/mL, gifted by the laboratory of Henning Walczak) for up to 48 h. Where multiple time points were used, stimulations were performed in a reverse time-course fashion so that all organoids were harvested at the same time. XELJANZ (tofacitinib; 20 µM, Pfizer), necrostatin-1s (10 μM, nec1s, Merck, 504297), GSK872 (10 μM, GSK872, SynKinase, SYN-5481), VX-765 (40 μM, Selleck, S2228), MCC950 (10 μM, kindly provided by A. Roberson and M. Cooper, University of Queensland, Australia), Q-VD-OPh (40 μM, QVD, MedChemExpress, HY-12305), Human Fas Ligand/TNFSF6 Antibody (10 μg/mL, R&D, MAB126), Human TRAIL/TNFSF10 Antibody (10 μg/mL, R&D, MAB375), ABT-737 (1 mM, Active Biochem; A-6044), cycloheximide (10 mg/mL, Sigma; C7698), TNF (50 ng/mL), Smac mimetic (1 mM, Compound A, TetraLogic Pharmaceuticals), Q-VD-OPh (20 μM), ABT-199 (1 mM, Active Biochem; A-1231), S63845 (10 mM, Active Biochem; A-6044) and A-1331852 (BCL-XL inhibitor; AbbVie, provided by Guillaume Lessene, WEHI) stimulations were performed as described in the relevant figure legends.

### CRISPR gene editing of organoids via ribonucleoprotein and electroporation

Human *NOS2* knockout intestinal organoids were generated using the Alt-R™ CRISPR-Cas9 system from Integrated DNA Technologies (IDT) and a published protocol (*108*). The Alt-R™ CRISPR-Cas9 crRNA (2 nM) was designed using IDT’s online software (https://sg.idtdna.com/site/order/designtool/index/CRISPR_PREDESIGN) The sequence is listed in Table S1. Alt-R® CRISPR-Cas9 tracrRNA, ATTO™ 550 (5 nM, IDT, 1075927) was resuspended and mixed to a final concentration of 50 µM, heated to 95°C for 5 minutes, then cooled to room temperature. To form the ribonucleoprotein (RNP) complex, Alt-R™ S.p. Cas9-GFP V3 (100 μg, IDT, 10008100) was diluted to 5 µg/µL and mixed with gRNA (50 µM) at a 1:2.5 molar ratio (3 µl sgRNA + 2.3 µl Cas9). The mixture was incubated at room temperature for 20 minutes.

Mature organoids (day 7) were harvested and digested into a single-cell suspension, with 1×10^6^ cells used for each electroporation reaction. Cells were centrifuged at 300 *g* for 5 minutes at 4°C, washed twice with PBS, and resuspended in 20 µL of P3 primary nucleofection solution (Lonza, V4XP-3032). The nucleofection mix consisted of 5.3 µL RNP complex, 1.2 µL IDT electroporation enhancer (1075915), and 20 µL cell/P3 nucleofection solution, for a total volume of 30 µL. The mixture was added to the supplied Nucleocuvette® Strip and transfected using the 4D-Nucleofector® Core Unit (Lonza, AAF-1003B) and 4D-NucleofectorTM X Unit (20 µL format) (Lonza, AAF-1003X) on program code DS-138.

Immediately after electroporation, cells were transferred to Eppendorf tubes, washed with ice-cold PBS, and resuspended in 500 µL FACS buffer containing DAPI (Thermo Fisher Scientific, 00-4959-52) for viability staining. The cell suspension was passed through a 40 µm filter and sorted by fluorescence-activated cell sorting (FACS) to isolate single live cells that were DAPI-negative and positive for both ATTO^TM^ 550 and GFP. Sorted cells were collected in Eppendorf tubes containing DMEM/F-12 with 1% BSA and 10 μM Y27632. Collected cells were washed and resuspended in fresh OGM with 5% matrix gel and 10 μM Y27632 before being returned to non-treated 24-well plates for expansion. Organoids were cultured for 7-10 days before harvesting for assessment of genome editing efficiency by immunoblot, as detailed in relevant figures and figure legends.

### Genetic modification of organoids using viral transduction

*BBC3* sgRNAs or scramble sgRNAs were designed and cloned into the lentiCRISPR v2 Cas9 backbone (a kind gift from F. Zhang, http://n2t.net/addgene:52961; RRID:Addgene_52961) (*109*). The sgRNAs (listed in Table S1) were introduced into intestinal organoids using lentiviral transduction. Lentiviral particles were generated based on a CRISPR/Cas9 protocol described previously (*110*). Briefly, HEK293T cells were plated at a density of 1×10^6^ cells per well in a 6-well plate. 1.48 μg of plasmid DNA and lentiviral packaging vectors (0.74 μg pMDL, 0.37 μg RSV-REV, 0.44 μg VSVg) were mixed with Lipofectamine 2000 (Thermo Fisher, 11668027) and OptiMEM and added to cells according to the manufacturers protocol. The mixture was incubated overnight then was replaced with DMEM containing 10% FCS the following day. Viral supernatants were collected after 48 hours, filtered through a 0.45 mm filter, and concentrated 20 times using a Lenti-X^TM^ Concentrator (Takara Bio, 631232) according to the manufacturer’s directions. The lentiviral particles were resuspended in OGM and aliquots stored at -80 °C. Transfected HEK293T cells were refreshed with DMEM containing 10% FCS medium, and the same procedure repeated 72 hours post-transfection.

Organoids cultured for seven days were digested into single cells for lentivirus transduction. Cells were resuspended in 250 μl OGM and seeded into non-treated 24-well plates at 2×10^5^ cells per well, then mixed with 250 μl of concentrated lentiviral particles supplemented with 8 μg/mL Polybrene (Sigma-Aldrich, TR-1003) and 10 μM Y27632. Four wells were used for each gene targeting experiment. The plate was centrifuged at 600 *g* for one hour at 32 °C, followed by incubation at 37°C with 5% CO_2_ for 6 hours. After incubation, the cells were collected, washed, and resuspended in fresh OGM with 5% matrix gel and 10 μM Y27632 before being returned to non-treated 24-well plates to allow organoids to grow. For puromycin selection, gene-modified organoids were digested into single cells and co-cultured with puromycin (2ug/mL) in OGM with 5% matrix gel and 10 μM Y27632 for 3 days. The surviving organoids were collected, washed, and resuspended in fresh OGM with 5% matrix gel and 10 μM Y27632 before being returned to non-treated 24-well plate to continue growth. Lentiviral particles containing doxycycline-inducible human BCL2 (cloned into pFRE3G PGK puro (*17*)) were generated in the same manner.

For constitutive overexpression of human BCL2, a retroviral pMIG BCL2 FLAG IRES GFP plasmid (gifted by the laboratory of John Silke) was used to generate retroviral particles by Lipofectamine 2000 co-transfection of HEK293T cells with retroviral packaging vectors (VSVg and gagpol), using 4 μg and 6 μg per well of a 10-cm plate. Virus supernatants were collected, filtered, and concentrated 20 times using Retro-X™ Concentrator (Takara Bio, 631455) according to the manufacturer’s directions and organoids infected with virus particles following the same procedure as for lentiviral transduction (see above). Cell sorting for GFP^+^ cells was used to enrich for organoids overexpressing BCL2. Briefly, infected organoids were digested into single cells and resuspend in FACS buffer (1% FCS in PBS) supplemented with propidium iodide (PI, 10 mg/mL) and 10 μM Y27632. Via fluorescence-activated cell sorting, single live cells (PI negative) positive for GFP were sorted into Eppendorf tubes containing DMEM/F-12 with 1% BSA and 10 μM Y27632. Collected cells were washed and resuspended in fresh OGM with 5% matrix gel and 10 μM Y27632 before being returned to non-treated 24-well plate for expansion.

### Database design

Quantitative and qualitative data from 5 arms of the study (patient history, clinical history, clinical data, histopathology scores, immunoblot quantitation) were managed using REDCap (*111, 112*) and Microsoft Excel. Due to the complexity of the RNA sequencing data, these data were instead managed separately using RStudio v2024.4.2.764. All data were linked to the patient’s study number (parent term) and the endoscopic classification of each biopsy (subparent term) to facilitate multiparametric analysis across the different arms of the study.

### Data visualization and statistical analysis

For all data displayed as graphs, the measure of center, variance, and the number of independent replicates is stipulated in the respective figure legend. The numbers represented in all heatmaps are provided in Table S1. Statistical tests were only applied to data that had been performed at least three time independently. The statistical test and the p-value cutoffs are stipulated in the respective figure legends. Statistical tests were performed with GraphPad Prims v10, except for differential gene expression analyses (see above for details). For data in Fig.4-6: each data point (symbols) depicted in graphs of organoids represents an independent biological replicate (*i.e.,* a different donor), and the number of times each experiment was repeated is detailed in the figure legends. All graphs of HepG2, MDA-MB-231 and BMDMs experiments display data points from experiments performed on the same cell line. Replicates from *in vitro* experiments were acquired either on different days or by different researchers with separate reagents and are displayed as the mean ± standard error of the mean (SEM).

### Endoscopic scoring of intestinal inflammation

The Simple Endoscopic Score for Crohn’s Disease (SES-CD) was used for patients with CD: 0-2 (remission), 2-6 (mild); 6-15 (moderate), > 15 (severe) (*113*). The Mayo Endoscopic Score was used for patients with UC: normal/inactive (0); mild (1); moderate (2); and severe (3) (*114*).

### Histology and Immunohistochemistry

For cell pellet: Trypsinized cells were centrifuged at 670 ×*g* for 3 min at room temperature. The supernatant was discarded, cell pellets resuspended in 10% (v/v) neutral buffered formalin, incubated for 15 minutes at room temperature, and centrifuged at 670 × g for 3 minutes at room temperature. Cell pellets were resuspended in 50 –70 μL of HistoGel (Epredia Cat#HG-4000-012) pre-warmed to 56 °C and then pipetted onto ice-cold glass coverslips to set. Set pellets were stored in 70% (v/v) ethanol until paraffin-embedding.

For tissue samples: biopsies were fixed in 10% v/v Neutral Buffered Formalin for 24–72 h, paraffin embedded, and sectioned for immunohistochemistry as described (*115*). Staining for caspase-8 and cleaved caspase-3 was performed as described (*115*). For staining iNOS, sections were treated with low pH Retrieval buffer (DAKO, K800521-2) at 97 °C for 30 minutes and stained with iNOS (R&D MAB9502, 1:200) on a Dako Omnis platform for 1 h and secondary antibody for 30 minutes followed by substrate chromogen (DAB) (DAKO, GV82511-2) treatment for 10 minutes and counterstaining with haematoxylin. Stained slides were scanned on an Olympus VS200 (objective: 20x, numerical aperture 0.8, media dry; software: Olympus VS200). Where higher resolution was required, slides were scanned on the Olympus VS200 using the 60x objective (numerical aperture 1.42, media oil). Downstream analyses used QuPath Software (v0.4.3) (*116*).

For organoids: organoids were harvested from 2-3 wells of a 96-well plate and transferred into Eppendorf tubes. After centrifugation at 10,000 rpm for 1 minute, the organoid pellet was gently resuspended in 1 mL PBS. To ensure removal of residual Matrigel, the organoid samples were kept on ice for 1 hour, followed by centrifugation at 10,000 rpm for 1 minute. The pellet was resuspended in pre-heated HistoGel at 55-60°C, pipetted gently, and placed onto a pre-cooled cover slip. After solidification at 4°C for 15 minutes, the gel domes were fixed in 10% formalin at 4°C overnight. The fixed domes were then transferred to cryomolds, ready for paraffin embedding. Subsequent histological or immunohistochemical analyses were performed on sectioned paraffin-embedded organoids with antibodies against LGR5 (clone OTI2A2, Thermo Fisher Scientific, TA503316, 1:300, described in (*106*)). For Intestinal Alkaline Phosphatase Polyclonal Antibody (ALPI) staining, sections were treated with low pH Retrieval buffer (DAKO, K800521-2) at 97 °C for 30 minutes and stained with ALPI (Catalog # PA5-22210, 1:200) on a Dako Omnis platform for 1 h and secondary antibody for 30 minutes followed by substrate chromogen (DAB) (DAKO, GV82511-2) treatment for 10 minutes and counterstaining with haematoxylin. Slides were scanned on the Olympus VS200 using the 60x objective (numerical aperture 1.42, media oil). The displayed immunohistochemistry images of intestinal tissues and organoids were processed using ImageJ (*117*), with brightness and contrast adjusted to a range of 0-235 and gamma set to 1.5.

### Histopathological scoring of intestinal inflammation

The Robarts Histopathology Index (RHI) was used to measure IBD activity within biopsies (*118*). Scoring was performed by one anatomical pathologist with gastrointestinal expertise based on haemtoxylin and eosin-stained slides sliced from formalin-fixed paraffin-embedded biopsies. Slides were de-identified as to disease state (control or IBD) and inflammatory state (non-inflamed, margin or inflamed), however the anatomical location of the biopsy was known. When a sample could not be scored due to high levels of inflammation and tissue abnormalities (e.g. the tissue was largely comprised of inflammatory neutrophilic exudate) a pseudo-score of 15 that likely underrepresents the extent of disease activity was assigned.

### Quantitation of iNOS expression via immunohistochemistry

Images were analyzed using QuPath (v0.4.3) (*116*). A region of interest was manually defined to cover the whole section. The brown DAB signal was then thresholded to isolate the iNOS-positive areas. The iNOS-positive area as a percentage of the whole section was then measured.

### Cell and tissue protein lysates

For human tissue lysates: biopsies were transferred from ice-cold dPBS into 0.4 mL of ice-cold RIPA buffer (10 mM Tris-HCl pH 8.0, 1 mM EGTA, 2 mM MgCl2, 0.5% v/v Triton X-100, 0.1% w/v sodium deoxycholate, 0.5% w/v SDS, and 90 mM NaCl) supplemented with 1x Protease and Phosphatase Inhibitor Cocktail (Cell Signaling Technology Cat#5872) and 100 U/mL Benzonase (Sigma-Aldrich Cat#E1014) and then homogenized with a stainless steel ball bearing in a Qiagen TissueLyzer II (30 Hz, 1 minute). For organoid lysates: ∼1000-2000 organoids were collected, washed twice with ice-cold PBS, and lysed in 60-120 μL of ice-cold RIPA buffer supplemented with cOmplete Protease Inhibitor Cocktail (Roche Biochemicals, 11697498001), phosphatase inhibitors (Merck, 4906837001) and 100 U/mL Benzonase (Sigma-Aldrich Cat#E1014). HT29 cells or HEK293Ts were lysed in RIPA buffer supplemented with cOmpleteTM Protease Inhibitor Cocktail, phosphatase inhibitors, and 100 U/mL Benzonase.

### Immunoblot

The protein concentration of lysates was measured using Pierce™ BCA Protein Assay Kits (Thermo Fisher Scientific, 23225) according to the manufacturer’s directions. For Fig. 2 and fig. S2, 20 μg of HT29 cell lysate and 40-50 μg tissue lysates were boiled for 10 minutes in Laemmli sample buffer (126 mM Tris-HCl, pH 8, 20% v/v glycerol, 4% w/v SDS, 0.02% w/v bromophenol blue, 5% v/v 2-mercaptoethanol) and fractionated by 4–12% Bis-Tris gel (Thermo Fisher Scientific Cat#NP0335BOX) using MES running buffer (Thermo Fisher Scientific Cat#NP000202). Note, patient lysates were always run alongside HT29 cell standards to allow the quantitative comparison of immunoblot data across this study. After transfer onto polyvinylidene fluoride (Merck Cat# IPVH00010), gels were Coomassie-stained as per manufacturer’s instructions (Thermo Fisher Scientific Cat#LC6060) and membranes were blocked in 5% w/v cow’s skim milk powder in TBS containing 0.1% Tween 20 (TBS+T) and then probed with primary antibodies (1:2000 dilution for rat primary antibodies or 1:1000 for other primary antibodies in blocking buffer supplemented with 0.01% w/v sodium azide; see Antibodies section for details) overnight at 4 °C, washed twice in TBS + T, probed with an appropriate HRP-conjugated secondary antibody (see Antibodies section for details), washed four times in TBS + T and signals revealed by enhanced chemiluminescence (Merck Cat#WBLUF0100) on a ChemiDoc Touch Imaging System (Bio-Rad). Between probing with primary antibodies from the same species, membranes were incubated in stripping buffer (200 mM glycine pH 2.9, 1% w/v SDS, 0.5 mM TCEP) for 30 minutes at room temperature and then re-blocked.

For all other immunoblot figures, protein concentration was adjusted into 1 mg/mL. Lysates were subsequently mixed with SDS-PAGE sample buffer. Proteins from lysates were separated using 4%–12% gradient gels (Invitrogen) and then transferred onto either nitrocellulose membranes (Amersham) or Immobilon-P polyvinylidene fluoride membranes (Merck Millipore; IEVH85R). To ensure accurate protein loading of organoid lysates, Ponceau staining was routinely performed. The membranes were blocked for 30 minutes with 5% skim milk (Devondale) in (TBS+T) at room temperature. After blocking, the membranes were incubated with primary antibodies overnight at 4°C. The primary antibodies were diluted in 5% BSA TBS+T with 0.04% sodium azide, typically at a dilution of 1:1000 unless otherwise specified. Between probing with primary antibodies from the same species, membranes were incubated in stripping buffer (200 mM glycine pH 2.9, 1% w/v SDS, 0.5 mM TCEP) for 30 minutes at room temperature and then re-blocked. See Antibodies section for details. Appropriate HRP-conjugated secondary antibodies (see Antibodies section for details) were diluted 1:5000∼10,000 in 5% skimmed milk in TBS+T and applied to the membranes for 1 hour at room temperature. The membranes were washed three times for 5 minutes each in TBS+T between antibody incubations, and four times for 5 minutes each after the secondary antibody incubation. The membranes were developed using ECL (Millipore, Bio-Rad) and visualized with the ChemiDoc Touch Imaging System (Bio-Rad) using Image Lab v6.1 (Bio-Rad).

### Immunoblot quantitation

For Fig. 2 and fig. S2, densitometric analysis of the raw full-resolution .scn Chemidoc files was performed using Image Lab v6.1 (Bio-Rad). For phosphorylation or cleavage events, their densitometric signals were expressed relative to their parent protein (e.g. phosphorylated RIPK3 values were expressed relative to non-phosphorylated RIPK3, and cleaved caspase-3 values were expressed relative to the total caspase-3 signal). For unmodified targets, their densitometric signals were first expressed relative the same unmodified protein in a HT29 cell standard and then adjusted for differences in GAPDH (e.g. cIAP2 levels in one biopsy lysate was expressed relative to cIAP2 levels in 20 μg of untreated HT29 cell lysate, and then further adjusted for differences in GAPDH between these lysates).

For fig. S5 and Fig. 6, densitometric analysis was performed using ImageJ v1.54h (*117*). Immunoblot images were first converted to grayscale (8-bit), and background noise was reduced by applying a background subtraction with a rolling ball radius of 50 pixels. The images were then inverted to prepare for band selection. For each band of interest, integrated density (IntDen) was quantified. For PUMA analysis, the integrated density values for each band were normalized to TNF/Compound A (TS) treated HT29 cells sample on the same membrane, then adjusted for differences in GAPDH. For iNOS analysis, the integrated density values for each band were normalized to non-IBD control on each membrane, then adjusted for differences in GAPDH.

### IncuCyte cell death measurements

To quantify organoid viability using the IncuCyte S3 (Sartorius) system, organoids were plated into non-treated 96-well plates as outlined above. IncuCyte® Cytotox Red Dye (Sartorius, 4632) was added to stain the dead cells at a dilution of 1: 20000 one hour before imaging. Throughout the assay, both brightfield and fluorescent images were collected with the phase/brightfield channel and red fluorescence channel (250ms exposure) using a 4× objective and Spheroid scan type. Images of both channels were exported and analysed through ImageJ 1.53t using a custom semi-automated macro. In brief, pairs of brightfield and red fluorescence images were cropped as shown in fig. S4D. The “MorphoLibJ” plugin (*119*) was used to preprocess the brightfield images, then the images converted into a binary mask based on a threshold range of 0 to 132 pixel values. The cropped “Regions of Interest (ROI)” was measured and defined as the total organoid area. For red fluorescence images, the positive area was measured within the brightfield ROI and based on the manual threshold mask. Cell death was assessed by the percentage of red fluorescent positive organoid area relative to the total organoid area. MDA-MB-231, HepG2 and BMDM viability was also measured through IncuCyte S3 (Sartorius) imaging. Cells were seeded in triplicate in 96-well plates and were incubated in SPY505-DNA (1:1000, Spirochrome, SC101) and propidium iodide (PI, 0.3 mg/mL, Sigma-Aldrich, P4170) for 3 hours before imaging. Throughout the assay, both phase and fluorescent images were collected using the Phase channel, Green fluorescence channel (300 ms exposure) and Red fluorescence channel (400 ms exposure) with a 10× objective and the Scan type of Standard. The algorithm setting was adjusted to accurately detect green and red puncta. To calculate the % PI positive cells (*i.e.,* cell death), the puncta count for the red channel was divided by the green channel count and the result multiplied by 100.

### Intestinal biopsy collection

Adults with or without IBD scheduled for endoscopic evaluation of the lower gastrointestinal tract (flexible sigmoidoscopy or colonoscopy) with the Gastroenterology Department at the Royal Melbourne Hospital were screened for eligibility. Patients were ineligible for recruitment if they had: active infection, active malignancy, simultaneously or currently received anti-tumor therapy, non-steroidal anti-inflammatory drug use in the last month, hereditary or familial polyposis syndromes, non-IBD forms of colitis (e.g. microscopic colitis, ischemic colitis, diversion colitis, or diverticulitis). Eligible patients were recruited and consented via a signed form. In total, 80 patients were recruited from 30 August 2021 to 21 April 2023 (see Table S1). One recruited patient was subsequently excluded from this study due to a finding of probable colorectal cancer during endoscopy, with total 80 patients included in our study. For patients with IBD, intestinal biopsies were retrieved endoscopically from relatively non-inflamed, marginally inflamed and inflamed areas of intestine. Where there were no signs of endoscopic inflammation, biopsy collection was instead tailored to the individual’s prior sites of disease activity (e.g. if a patient with ileal CD was found to have a SES-CD of 0, then the ileum was deemed to be the “inflamed” region with distal parts of the bowel deemed to be the “margin” and “non-inflamed” sites). Patients without IBD (non-IBD controls), had biopsies retrieved endoscopically from only non-inflamed segments of the intestine, with comparable segments sampled in each patient. Boston Scientific Radial Jaw^TM^ biopsy forceps and Olympus EVIS EXERA III endoscopes were used for biopsy collection. Upon retrieval, matched biopsies were immediately placed into the following ice-cold media: 1) 10% v/v neutral buffered formalin, 2) dPBS (Thermo Fisher Scientific Cat#14190144) supplemented with protease inhibitors (ThermoFisher Scientific Cat# A32955) and phosphatase inhibitors (ThermoFisher Scientific; Cat#A32957), 3) RNAlater (ThermoFisher Scientific; Cat#AM7021), 4) 95% v/v methanol and 5% v/v acetic acid, and 5) DMEM/F-12 with 15 mM HEPES (DMEM/F-12, STEMCELL Technologies, 36254) and 1% w/v Bovine Serum Albumin (BSA, Sigma-Aldrich, A4612). Two biopsies were collected per site/patient/media.

Four organoid lines were prepared from normal human large intestine tissue samples (WCB316, 312, 317, 308). Samples were collected from resection margins of patients with colorectal cancer undergoing surgery from the Royal Melbourne Hospital, Western Health, Eastern Health and the Northern Hospital in Melbourne, Australia, between November 1, 2021 and March 31, 2022. All patients provided written informed consent, and the study was approved by medical ethics committees at all sites (HREC 2016.249, WEHI).

### Clinical scoring of disease activity

Patients were interviewed prior to endoscopy for contemporaneous clinical symptomology. Patients with CD were scored using the Harvey Bradshaw Index (HBI): remission 0-5; mild 5-7; moderate 8-16; severe >16 (*120*). Patients with UC were scored using the Simple Clinical Colitis Activity Index (SCCAI): remission </= 2; mild 3-5; moderate 6-11; >12 severe (*121*).

### Nitric oxide production Griess assay

Griess assays were performed as previously described (*53*). Cell supernatants were assayed in duplicate alongside a sodium nitrite standard curve ranging from 100 µM to 1.56 µM in OGM or ODM, which was assayed in triplicate. To each 25 µL of cell supernatant, 25 µL of sulfanilamide (1% w/v, Sigma) in phosphoric acid (5% v/v, Sigma) was added and incubated for 5 minutes. This was followed by the addition of 25 µL of N-1-napthylethylenediamine dihydrochloride (0.1% w/v, Sigma) in water. Absorbance was measured using a CLARIOstar Plus Microplate Reader (BMG LABTECH) and then interpolated from the background-corrected sodium nitrite standard curve.

### Caspase DEVDase activity assay

Organoids were collected and lysed with DISC buffer (20 mM Tris-HCL pH7.5, 150 mM NaCl, 1% Triton X-100, 2 mM EDTA and 10% Glycerol) supplemented with cOmpleteTM Protease Inhibitor Cocktail and phosphatase inhibitors. A substrate mixture of Ac-DEVD-AMC substrate (20 µM, BD Pharmingen, 556449), dithiothreitol (2 mM) and Protease Assay Buffer (20 mM HEPES pH 7.5, 10% glycerol) was made up fresh immediately before use. 20 µL of cell lysates were incubated with 200 µL of substrate mixture at 37 °C for 1 hour. Fluorescence was measured using a using a CLARIOstar Plus Microplate Reader (BMG LABTECH) with an excitation wavelength of 380 nm and an emission wavelength range of 430-460 nm.

### Research ethics

Ethical approval for intestinal tissue collection from participants undergoing endoscopy procedures through the Gastroenterology Department at the Royal Melbourne Hospital was attained from the Human Research Ethics Committee (HREC): HREC 2021.074. This was in accordance with the National Health and Medical Research Council (NHMRC) National Statement on Ethical Conduct in Human Research (2007) and the Note for Guidance on Good Clinical Practice (CPMP/ICH-135/95). Site-specific governance was sought for each of the collaborating sites: WEHI and the University of Melbourne. Collaboration amongst all three participating institutions was officiated through the Melbourne Academic Centre for Health Research Collaboration Agreement (Non-Commercial). The human research in this study was performed in accordance with the principles expressed in the World Medical Association Declaration of Helsinki and conforms to the principles set out in the United States Department of Health and Human Services Belmont Report. The human materials used in this study were obtained with signed informed consent from all subjects.

### RNA isolation and quantitative polymerase chain reaction (qPCR) from organoids

Total RNA was isolated from organoids using the ISOLATE II RNA Mini Kit (Bioline, 52073) following the manufacturer’s guidelines. cDNA was reverse transcribed from 1 μg of RNA using the SuperScript III Reverse Transcriptase (Invitrogen, 18080-085), oligo (dT) nucleotides (Promega, C110B-C). Quantitative Real-Time PCR (qRT-PCR) was performed on cDNA samples or nuclease-free water (control) using Maxima SYBR Green/ROX qPCR Master Mix (Thermo Fisher Scientific, K0223) and the ViiA 7 Real-time PCR system (Applied Biosystems). Samples were run in duplicates. Relative gene expression was normalized to the housekeeping reference gene *HPRT* and presented as the fold-change relative to unstimulated control sample or OGM sample, analysed using the ΔΔCt method (*122*). qRT-PCR primer sequences used in this study are listed in Table S1.

### Single-cell RNA sequencing data re-analysis

The raw counts of ‘Non-IBD ’, ‘IBD non-inflamed’ and ‘IBD inflamed’ samples from (*123*) were quality controlled using an R package scater (*124*). To analyse and visualize the high-dimensional scRNA-seq data, we employed Principal Component Analysis (PCA) and Uniform Manifold Approximation and Projection (UMAP). The ‘IBD inflamed’ data were subset and renormalized using the same pipeline. A gene-set score for each individual cell in the sub-dataset was calculated using AUCell (*125*) based on the cluster B marker list (Table S1), respectively. The differential gene expression analysis was performed using a pseudo-bulked approach by mastR (*97*), which aggregated the raw counts into each pseudo-bulked samples by ‘state’, ‘Sample’ and ‘ident’. The pseudo samples with less than 20 cells were excluded, leaving in 62 samples for analysis.

## Acknowledgments

We thank the Royal Melbourne Hospital Endoscopy Unit (3W) and Anaesthetics staff, Andrew Trinh, Eva Zhang, James Winston, Siddharth Sood, Allie Mack, Peter Tagkalidis and Peter Prichard for their assistance with endoscopic biopsy retrieval. We are grateful to patients for donating their tissues for this study. We thank Sebastian Hughes for assistance with the immunoblotting of patient samples. We thank Anne Hempel (WEHI), Bruce Rosengarten (Crohn’s & Colitis Australia) and Ian Wicks (WEHI) for constructive feedback of this work. We thank WEHI Monoclonal antibody facility for producing several antibodies used in this study. We thank the WEHI Histology team for high-level support with immunohistochemistry. We acknowledge the use of BioRender with the preparation of figures.

## Author contributions

Conceptualization: JMM, ALS, AHA, BC, JEV, JP. Methodology: JMM, ALS, AHA, BC, JEV, JP. Investigation: AHA, JP, KMP, SNY, IK, MB, JAR, WC, AVJ, AJ, CRH, SC, TT, WL, SS, ALS. Analysis: AHA, JP, IK, SF, JC, AW, LWW, PR, IA, TS, ALS. Resources: AHA, SS, AM, AP, NS, GI, DS, AE, WB, FMc, BC, GL, HW, OMS, ALS; Supervision: YZ, OS, LGR, EDH, KLR, RB, SEN, KEL, BC, ALS, JEV, JMM. Writing – original draft: JMM, ALS, AHA, BC, JEV, JP. Writing – review & editing: all authors.

## Data and materials availability

Full resolution raw blot data will be uploaded to FigShare upon paper acceptance. The bulk RNA sequencing data will be uploaded to the Gene Expression Omnibus repository upon paper acceptance.

## Figures & Legends

**Fig. S1.**
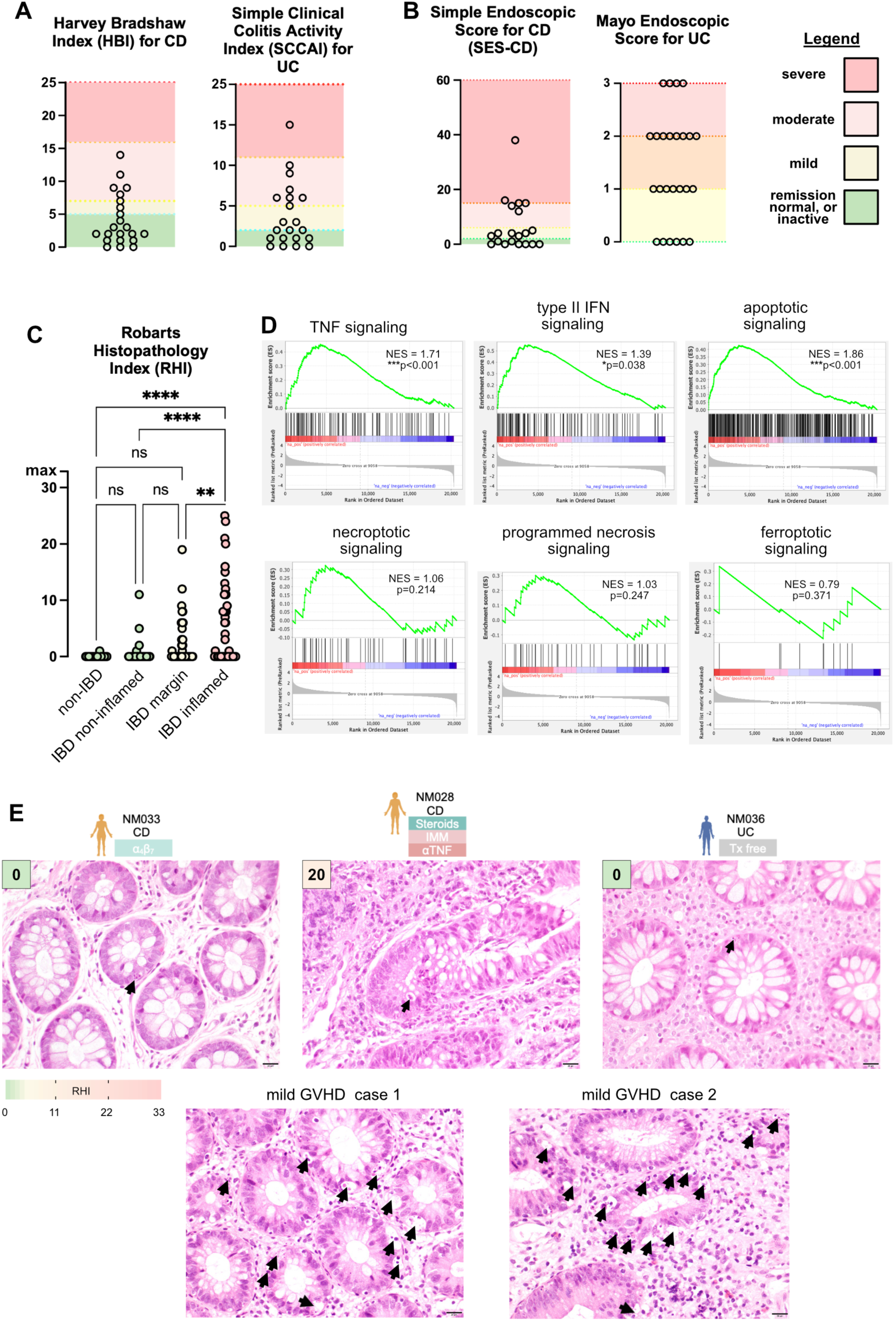
Cell death is not a histologically prominent feature of IBD. **(A-B)** Clinical scores (A) and endoscopic scores (B) for the patients with CD and UC included in this study. Each dot represents one patient. **(C)** Blinded histopathology scores (RHI) for all biopsies collected for this study. Each dot represents one biopsy. ****p<0.0001 by one-way ANOVA. **(D)** Gene set enrichment plots for the stipulated inflammatory and cell death signaling pathways. **(E)** Hematoxylin and eosin-stained sections of intestinal biopsies from patients with CD, UC or graft-versus-host disease (GVHD). Representative micrographs were from biopsies where subsequent immunoblots indicated the presence of caspase-3 cleavage. Black arrows indicate apoptotic bodies that were annotated in a blinded manner by a histopathologist. The study number and treatment of the patients with IBD is shown. Boxed numbers indicate the histopathological score (RHI) of each biopsy site. Scale bars are 20μm.

**Fig. S2.**
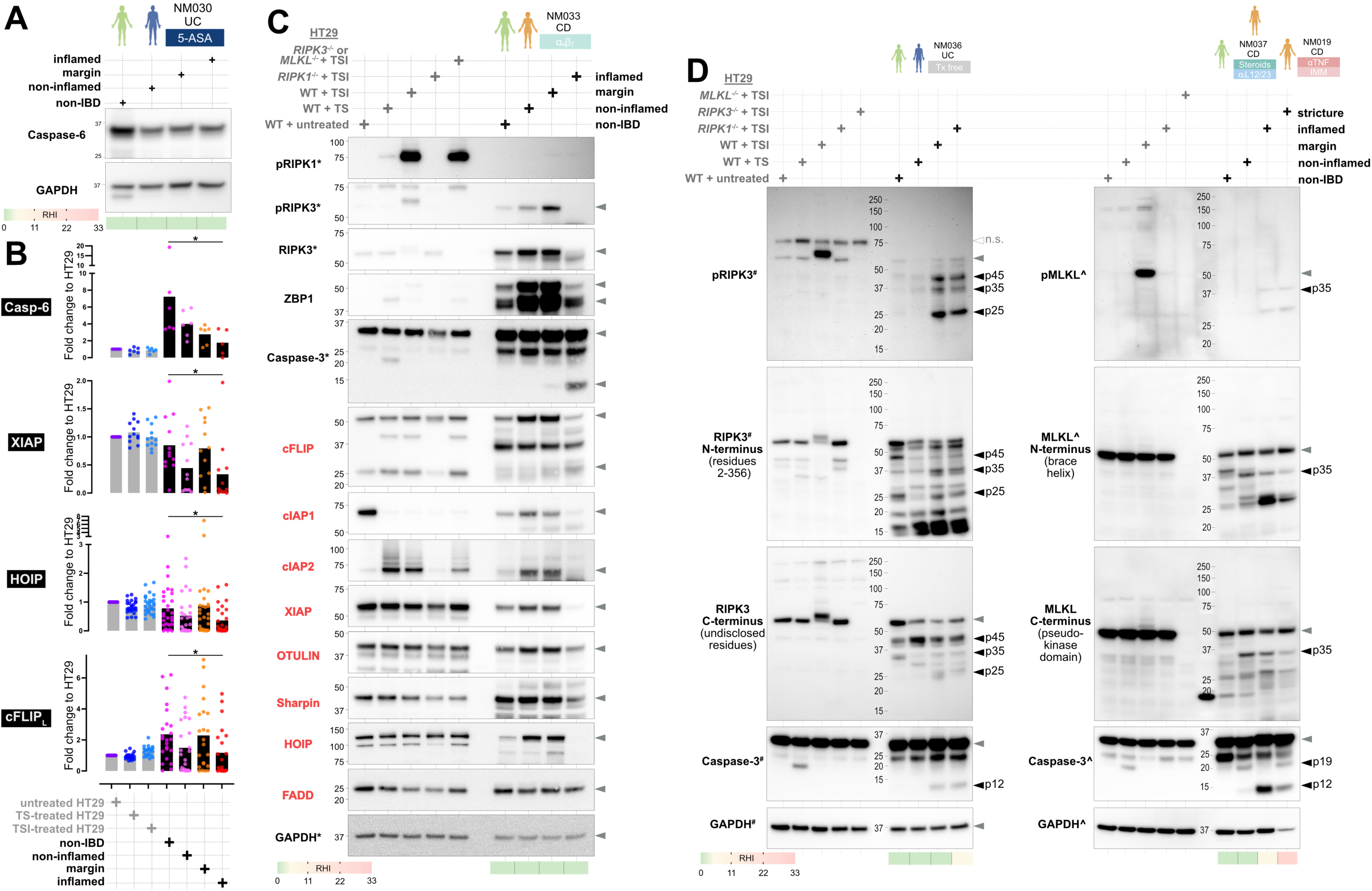
Cell death disinhibition coincides with apoptosis and RIPK3/MLKL cleavage. **(A)** Immunoblot of intestinal biopsies from patients. The study number (NM), IBD subtype (UC/CD) and treatment of the patient is stipulated. The histopathological score (RHI) of each biopsy site is shown. **(B)** Graphs show relative expression levels of the indicated proteins in HT29 cell and biopsies. Each dot represents one biopsy. Bars indicate mean values. *p<0.05 by one-way ANOVA with Geisser-Greenhouse correction. **(C-D)** Immunoblot of lysates from HT29 cells (grey text) and intestinal biopsies from patients. Non-specific (n.s.). Immunoblot data annotated with an asterisk (*) in panel C were previously published in Fig. EV4 of (*41*). Immunoblot data annotated with a hastag (#) in panel D also appeared in Fig. 2A of this manuscript. Immunoblot data annotated with a caret (^) in panel D also appeared in Fig. 6C of (*41*).

**Fig. S3.**
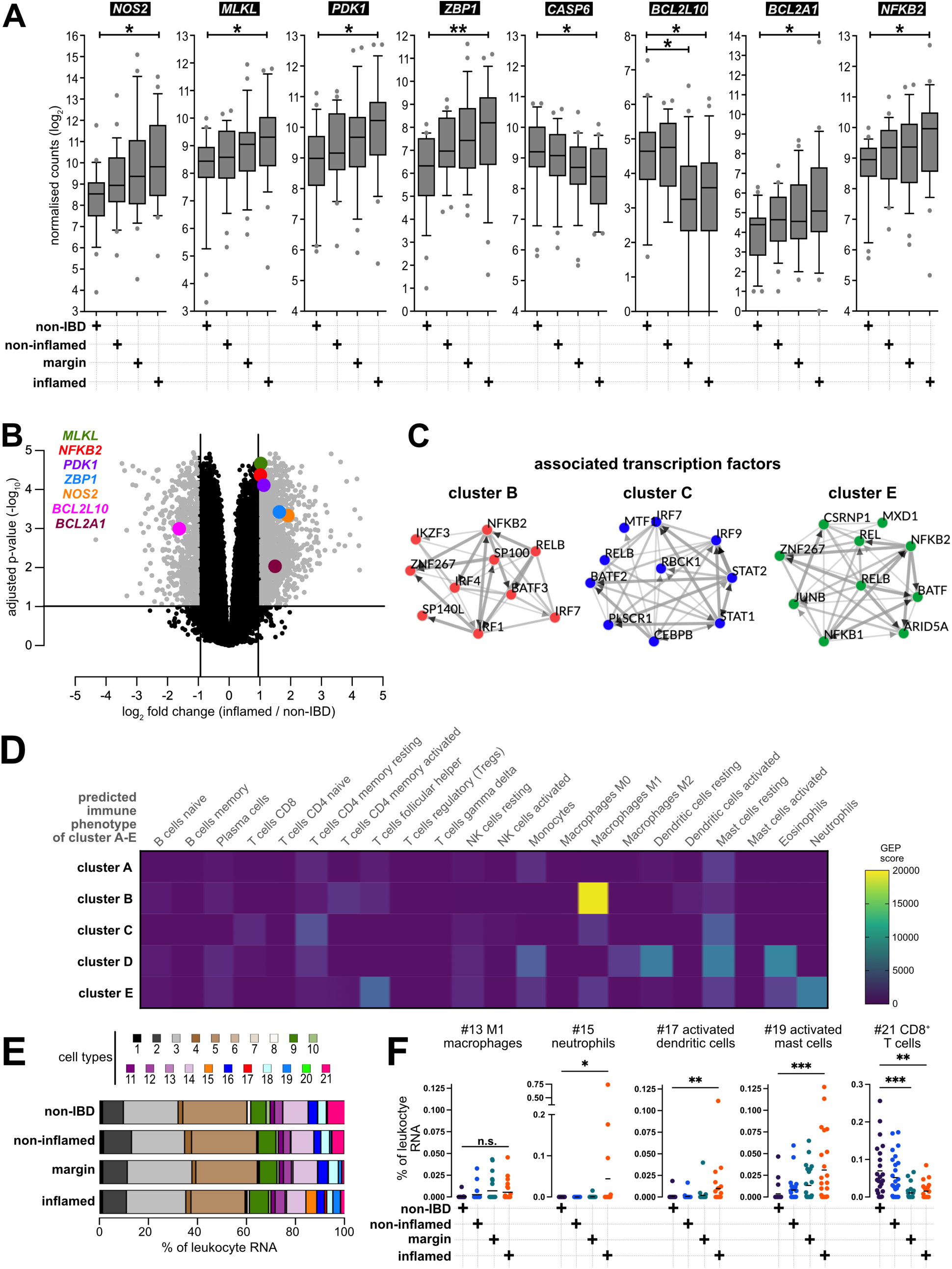
The profile of inflamed epithelia resembles an anti-pathogen M1-macrophage response. **(A)** Transcript levels from bulk RNA sequencing of intestinal biopsies from patients with IBD (n=26 non-inflamed, n=24 margin, n=25 inflamed biopsies) and non-IBD patients (n=23). Plots indicate 10^th^, 50^th^ and 90^th^ percentiles with dots showing values beyond this range. *p<0.05 and **p<0.01 by one-way ANOVA with Kruskal-Wallis correction. **(B)** Plot shows the differential expression of transcripts between inflamed IBD (n=25) and non-IBD (n=23) samples. Adjusted p-values determined by limma-voom pipeline (see Methods). **(C)** The top 10 transcription factors that regulate the expression of genes in clusters B, C and E as identified by (*89*). The thickness of the arrow is proportional to the number of sources supporting an interaction between transcription factors. **(D)** Bulk RNA sequencing data from all intestinal samples (n=98; n=23 non-IBD, n=75 IBD) were subjected to correlation analysis between the gene expression profiles (GEP) of immune subsets and clusters A-E. **(E-F)** Deconvolution of immune subsets from the bulk RNA sequencing of intestinal biopsies (n=26 non-inflamed, n=24 margin, n=25 inflamed, n=23 non-IBD). Cell type-specific reads were expressed as a percentage of total leukocyte reads per group in panel E, or per biopsy in panel F. One dot per biopsy. *p<0.05, **p<0.01, ***p<0.001 by one-way ANOVA with Kruskal-Wallis correction. Cell types: 1) naïve B-cell, 2) memory B-cell, 3) plasma cell, 4) naïve CD4^+^ T-cell, 5) resting memory CD4^+^ T-cell, 6) activated memory CD4^+^ T-cell, 7) follicular helper T-cell, 8) regulatory T-cell, 9) resting NK-cell, 10) activated NK-cell, 11) monocyte, 12) M0-macrophage, 13) M1-macrophage, 14) M2-macrophage, 15) neutrophil, 16) resting dendritic cell, 17) activated dendritic cell, 18) resting mast cell, 19) activated mast cell, 20) eosinophil, 21) CD8^+^ T-cell.

**Fig. S4.**
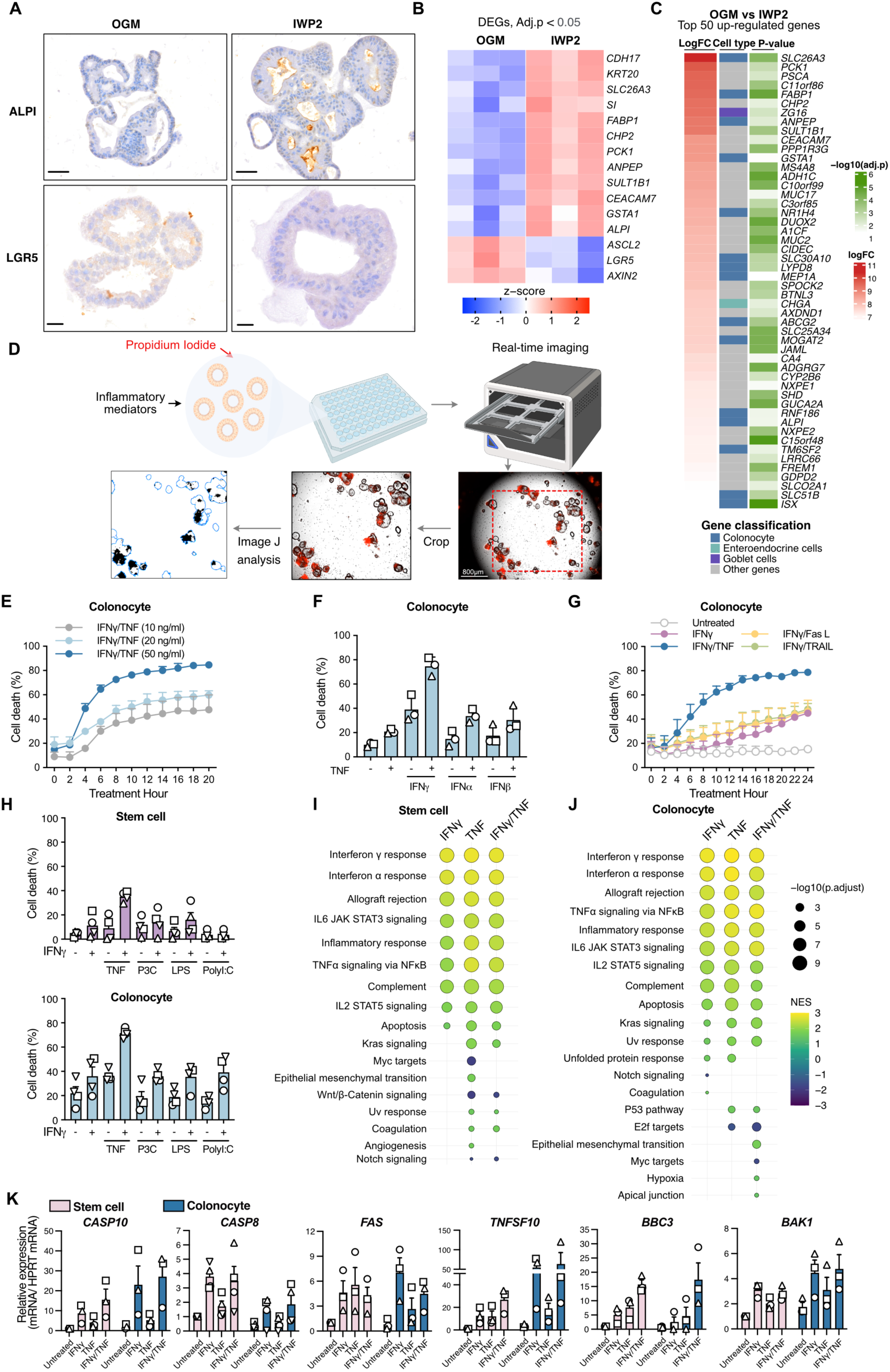
Impact of inflammatory cytokines and Toll-like receptor ligands on intestinal organoids. **(A**) Immunohistochemistry for LGR5 and ALPI in undifferentiated organoids (OGM) and colonocyte-enriched organoids (IWP2) from one representative patient. Scale bars are 50 µm (top) and 20 µm (bottom). **(B)** Heatmap shows the differential gene expression of cell type markers between undifferentiated organoids (OGM) and colonocyte-enriched organoids (IWP2). n = 3 different donors. All entries with an adjusted p<0.05 by empirical Bayes moderated t-statistic and Benjamini–Hochberg multiple test correction. **(C)** Heatmap of top 50 upregulated genes in IWP2 differentiated organoids compared to OGM organoids. The middle column highlights genes that belong to GI tract epithelial cell type markers (PanglaoDB). **(D)** Schematic of IncuCyte real-time imaging used to quantify intestinal organoid cell death by Cytotox Red Dye staining in response to inflammatory mediators. **(E)** Analysis of cell death in colonocyte organoids treated with indicated concentrations of IFNγ and TNF (n=3 different donors, mean ± SEM, pooled from 2 independent experiments). **(F)** Cell death in colonocyte organoids were treated with IFNγ (50 ng/mL), IFNα (100 ng/mL), or IFNβ (100 U/mL) with TNF (50 ng/mL) for 24 hours (n = 3 different donors, represented by different symbols). Data show the mean ± SEM, pooled from 2 independent experiments. **(G)** Cell death in colonocyte organoids treated with FAS ligand (10 ng/mL), TRAIL (5 ng/mL), TNF (50 ng/mL) and IFNγ (50 ng/mL) (n=4 different donors, mean ± SEM, pooled from 3 independent experiments). **(H)** Cell death (%) in stem cell or colonocyte organoids treated with TNF (50 ng/mL), LPS (50 ng/mL), Pam_3_CSK_4_ (P3C, 500 ng/mL) or PolyI:C (10 µg/mL) with IFNγ (50 ng/mL) for 24 hours (n=4 donors, represented by different symbols, mean ± SEM, pooled from 2 independent experiments). **(I, J)** Gene set enrichment analysis of organoids bulk RNA sequencing data (compare to untreated organoids). Normalized enrichment score (NES) and the nominal p-value are shown. **(K)** Relative gene expression levels assessed by qPCR in stem cell organoids and colonocyte organoids treated with IFNγ (50 ng/mL), TNF (50 ng/mL), or a combination of IFNγ and TNF for 3.5 hours (mean ± SEM, n = 4 donors for CASP8, n=3 donors for all others).

**Fig. S5.**
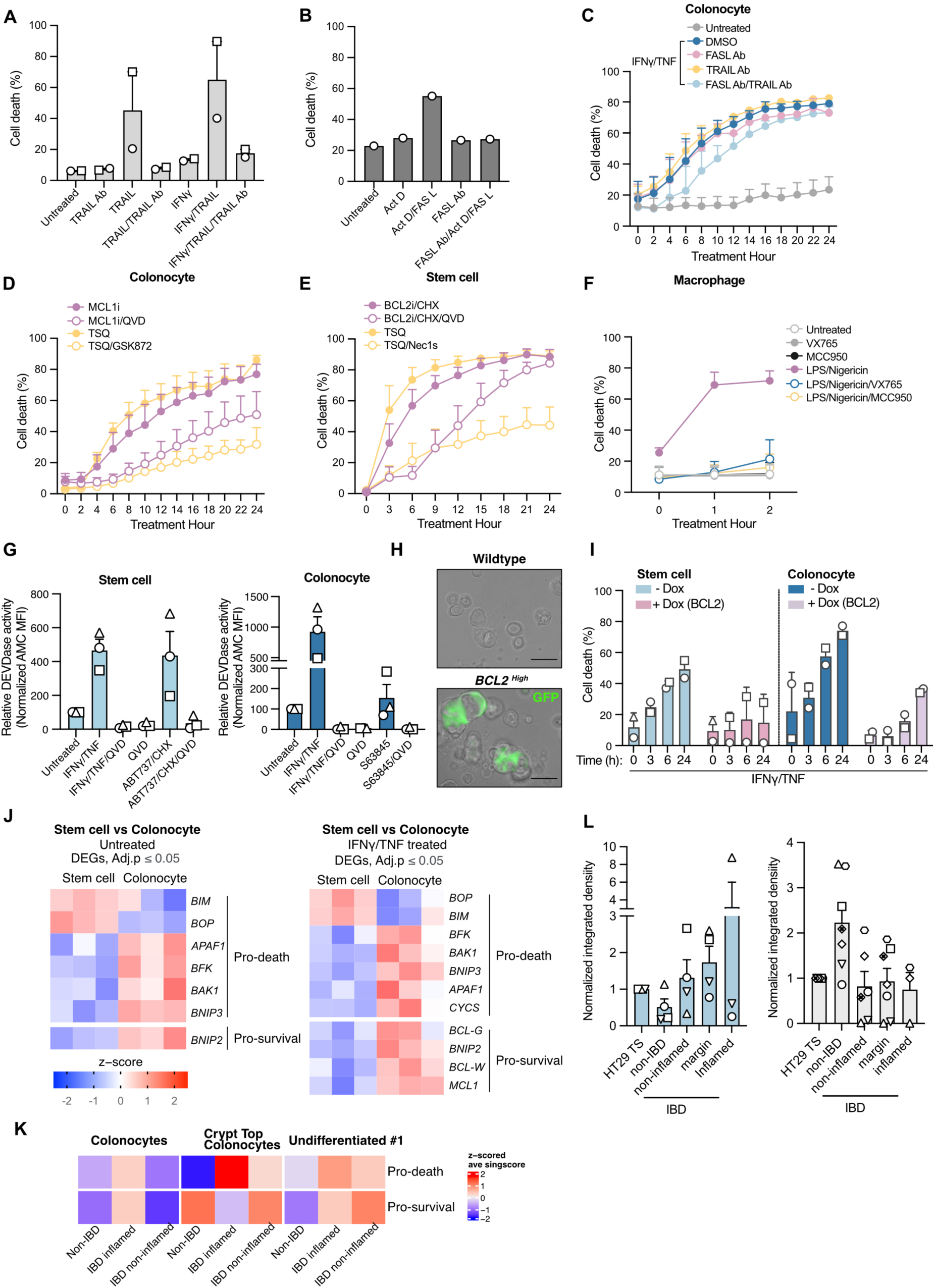
Fas and TRAIL neutralization does not block IFNγ/TNF killing of human intestinal organoids. **(A)** MDA-MB-231 cells were treated with IFNγ (50 ng/mL), TRAIL neutralizing antibody (TRAIL Ab, 10 μg/mL) and TRAIL (50 ng/mL) for 48 hours. Cell death was assessed by PI exclusion, measured by Incucyte® imaging (n = 2 independent experiments, mean ± SEM). **(B)** Cell death of HepG2 cells treated with TRAIL/TNFSF10 neutralizing antibody (FASL Ab,10 μg/mL) then Actinomycin D (Act D, 0.5 μg/mL) and FAS ligand (FasL, 100 ng/mL) was assessed by PI exclusion using IncuCyte imaging (n=1). **(C)** Cell death in colonocyte organoids pre-treated with vehicle (DMSO), TRAIL Ab (10 μg/mL), or FASL Ab (10 μg/mL) then IFNγ and TNF (n=3 different donors, mean ± SEM, pooled from 3 independent experiments). **(D, E)** Organoids were pre-treated with Q-VD-OPh (QVD, 10 μM), GSK’872 (10 μM), necrostatin-1s (Nec1s, 10 μM) or vehicle (DMSO) followed by stimulation with ABT-737 (1 μM) and cycloheximide (CHX, 10 mg/mL), or the MCL-1 inhibitor S63856 (5 μM) to activate mitochondrial apoptosis. Alternatively, intestinal organoids were stimulated with TNF(50 ng/mL), Smac-mimetic Compound A (1 μM) and QVD-OPh to induce necroptosis (TSQ). Cell death was assessed using Cytox Red uptake by IncuCyte live cell imaging (n=3 different donors, mean ± SEM, pooled from 3 independent experiments). **(F)** Mouse bone marrow-derived macrophages were primed with LPS (100 ng/mL) and treated with Nigericin (20 μM) ±pretreatment with VX-765 (20 μM) or MCC950 (10 μM) (n=3 different mice, mean ± SEM, pooled from 2 independent experiments. (**G)** Caspase-3 activity was determined by DEVDase activity assay in organoids (n=3 different donors, mean ± SEM, pooled from 3 independent experiments). **(H)** Representative immunofluorescence images of wildtype control and *BCL2* overexpressing intestinal stem cell organoids. Scale bar, 100 µm. **(I)** IncuCyte imaging was used to quantify cell death following induction of exogenous BCL2 by doxycycline (dox) in stem cell or colonocyte organoids and subsequent IFNγ and TNF stimulation. (n=2 donors, mean ± SEM, pooled from 2 independent experiments). **(J)** Heatmap of pro-survival and pro-death DEGs (see Figure 5E) comparing IFNγ/TNF treated stem cell organoids versus colonocyte organoids, or untreated stem cell organoids versus colonocyte organoids. All entries with an adjusted p<0.05 by empirical Bayes moderated t-statistic and Benjamini–Hochberg multiple test correction. **(K)** Single-cell IBD patient sequencing data (inflamed and non-inflamed mucosa from 3 non-IBD controls and 3 UC patients; (*52*)) was re-analyzed to define pro-survival and pro-death genes enrichment scores. **(L)** PUMA levels of non-inflamed, inflamed and marginal IBD tissues and non-IBD controls were analyzed by immunoblot and quantified by ImageJ (mean ± SEM); data represented separately as upregulated (n=4) and downregulated (n=7) trends. Protein levels are represented as the integrated band density and normalized to TNF/Compound A treated HT29 cells. PUMA levels of HT29 cells and biopsies are analyzed by immunoblot and quantified by ImageJ (mean ± SEM); Each dot represents one biopsy, data represented separately as upregulated (n=5, left panel) and downregulated (n=7, right panel) trends. Expression was represented as the integrated band density and normalized to TNF/Compound A treated HT29 cells then adjusted for differences in GAPDH.

**Fig. S6.**
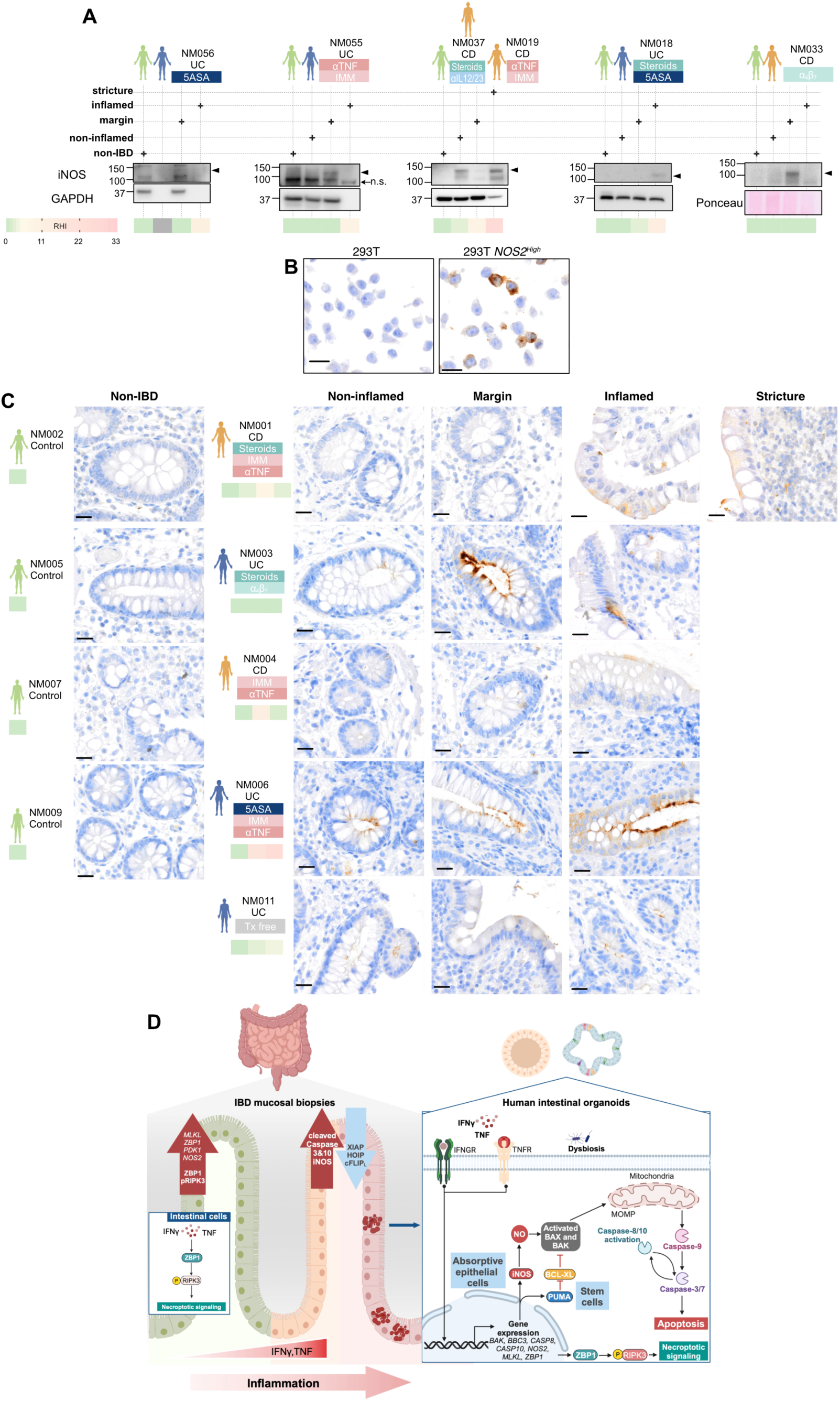
Increased epithelial iNOS levels in IBD patients. **(A)** Immunoblots showing iNOS levels in intestinal biopsies. The patient study number (NM), IBD subtype (UC/CD), treatment, and the histopathological score (RHI) of each biopsy site is shown. **(B)** 293T cells engineered to overexpress NOS2 and control 293T cells that do not express NOS2 were used to optimize and demonstrate NOS2 antibody staining specificity. Scale bar, 20 µm. **(C)** Representative immunohistochemistry for iNOS in intestinal tissue biopsies from n=5 patients with IBD and n=4 non-IBD patients. Scale bar, 20 µm. **(D)** Model depicting cell death signaling in IBD intestinal epithelium. Necroptotic signaling characterized by increased pRIPK3 and ZBP1 is prominent in histologically non-inflamed IBD tissue. In contrast, apoptotic signaling defined by elevated cleaved caspase-3/10 and downregulation of negative death regulators (HOIP, cFLIPL, and XIAP) is intensified in inflamed IBD tissue. The cell death signals correlate with altered TNF and IFNγ pathway expression. iNOS expression is also upregulated in IBD biopsies. Treating human derived intestinal organoids with TNF and IFNγ recapitulates IBD intestinal epithelium transcriptomically, triggers pro-apoptotic and pro-necroptotic gene expression as well as *NOS2* expression and cell death. Mechanically, iNOS expression activates BAX and BAK likely via the activity of nitric oxide (NO) in absorptive epithelial cells, while PUMA expression in intestinal stem cells inhibits BCL-XL to activate BAX and BAK. Both pathways end up with mitochondrial outer membrane permeabilization (MOMP) and cytochrome c release into the cytosol. Cytochrome c release and apoptosome formation promote caspase-3/7 activation, which further activates caspase-8/10 in a positive feedback loop, culminating in apoptosis.

